# Demography and linked selection interact to shape the genomic landscape of codistributed woodpeckers during the Ice Age

**DOI:** 10.1101/2022.02.04.479011

**Authors:** Lucas R. Moreira, John Klicka, Brian Tilston Smith

## Abstract

The glacial cycles of the Pleistocene had a global impact on the evolution of species. Although the influence of genetic drift on population genetic dynamics is well understood, the role of selection in shaping patterns of genomic variation during these dramatic climatic changes is less clear. We used whole genome resequencing data to investigate the interplay between demography and natural selection and their influence on the genomic landscape of Downy and Hairy Woodpecker, species co-distributed in previously glaciated North America. Our results revealed a dynamic population history with repeated cycles of bottleneck and expansion, and genetic structure associated with glacial refugia. Levels of nucleotide diversity varied substantially along the genomes of Downy and Hairy Woodpecker, but this variation was highly correlated between the two species, suggesting the presence of conserved genomic features. Nucleotide diversity in both species was positively correlated with recombination rate and negatively correlated with gene density, suggesting that linked selection played a role in reducing diversity in regions of low recombination and high density of targets of selection. Despite strong temporal fluctuations in N_e_, our demographic analyses indicate that Downy and Hairy Woodpecker were able to maintain relatively large effective population sizes during glaciations, which might have favored natural selection. The magnitude of the effect of linked selection seems to have been modulated by the individual demographic trajectory of populations and species, such that purifying selection has been more efficient in removing deleterious alleles in Hairy Woodpecker owing to its larger long-term N_e_. These results highlight that while drift captures the expected signature of contracting and expanding populations during climatic perturbations, the interaction of multiple processes produces a predictable and highly heterogeneous genomic landscape.

## Introduction

Pleistocene glacial cycles altered the distribution and evolution of entire communities [1,2]. Despite the profound impact glaciations had on the evolutionary trajectory of species, the majority of research on the topic has focused on how demographic dynamics have shaped neutral genetic variation [2,3]. Population expansion [4,5], genetic structuring in refugia [6–10], and decreased diversity in expanding populations [11–13] are among the most common patterns recovered. However, as species rapidly expanded and colonized areas under extreme environmental change they would have been subject to strong selective pressures, such as increased tolerance to cold and selection against deleterious mutations [14,15]. Understanding how natural selection, along with genetic drift, interact with features of the genome to shape the genomic landscape of diversity and differentiation will clarify the broader significance of the Ice Age on the evolution of species.

Demography and natural selection play a central role shaping levels of genetic diversity, but their effects are intertwined [16–18]. Neutral genetic diversity (θ) is the product of the rate at which new alleles are generated (i.e., mutation rate μ) by the effective population size (N_e_), so that diversity levels are predicted to increase as a function of the size of populations (In diploids, θ = 4N_e_μ [19,20]). On the other hand, fixation of beneficial alleles (selective sweep [21,22]) or removal of deleterious mutations (background selection [22–25]) can cause genetic diversity to decrease across the genome through the effect of linked selection [24]. Demographic perturbations that cause N_e_ to fluctuate over time and space (e.g., glacial bottlenecks) are, therefore, expected to result in a larger accumulation of mildly deleterious alleles when compared to large populations with constant N_e_ because of the reduced efficacy of purifying selection when genetic drift is strong [26–30]. Hence, populations resulting from founder events, such as at the leading edge of a postglacial expansion, often show elevated genetic load [27,30,31].

Levels of diversity and differentiation along the genome also vary due to the differing effects of intrinsic genomic properties [32–36]. Genome features such as variation in mutation rate, recombination rate, distribution of functional elements, and nucleotide composition impact the rates at which genetic variants are produced, maintained, and lost [37]. Regions enriched for functional elements (e.g., coding sequences), for instance, tend to exhibit significantly lower levels of genetic diversity due to the recurrent effect of natural selection [33,38–40]. The loss of variation is further amplified by linkage disequilibrium (LD), which reduces diversity at neutrally-evolving sites in close proximity to the targets of selection (hitchhiking effect [21]). The extent to which linked selection affects neighboring sites depends on the recombination rate, which shows considerable genome-wide variation [41–44]. Larger reductions in nucleotide diversity are expected to occur in genomic regions enriched for functional elements and with lower recombination rates. A correlation between nucleotide diversity, gene density, and recombination rate is therefore indicative that linked selection is at play. Quantifying covariance between evolutionarily independent species can help understand the interplay between these various conserved features of the genome and their impact on patterns of diversity and differentiation along the genome.

We aim to address drift-selection dynamics during the Pleistocene climatic cycles by estimating the impact of demography and linked selection on the genome of Downy (*Dryobates pubescens*) and Hairy (*D. villosus*) Woodpeckers, two co-distributed species that share similar ecologies and evolutionary histories. Downy and Hairy Woodpecker are year-round residents of a variety of habitats in North America, occurring in sympatry across an exceptionally broad geographic area from Alaska to Florida, although the range of the Hairy Woodpecker extends further south, reaching portions of Central America and the Bahamas [45]. Despite looking very similar, the two species are not sisters and share a common ancestor more than eight million years ago, without any evidence of recent hybridization [46,47]. During the glacial cycles of the Pleistocene, especially when the polar ice sheets reached their maximum extent (Last Glacial Maximum; 21 kya), a large portion of the present-day distribution of Downy and Hairy Woodpeckers were covered in ice, and populations of both species were restricted to southern refugia [12,48,49]. After the retreat of Pleistocene glaciers, Downy and Hairy Woodpeckers extended their distributions north, recolonizing higher latitudes. Phylogeographical studies in Downy and Hairy Woodpecker revealed that populations currently inhabiting previously glaciated areas show strong signatures of population expansion and population structuring consistent with multiple glacial refugia [12,48–51]. This shared demographic history provides an opportunity to investigate multiple genomic factors that might have impacted the distribution of diversity across populations and within the genomes of these two natural evolutionary replicates.

In this study, we generated whole-genome resequencing data for Downy and Hairy Woodpeckers to test whether the heterogeneous genomic landscape of diversity and differentiation in both taxa is correlated with intrinsic features of the genome, such as recombination rate and gene density, and whether differences in demographic history had an impact on the efficacy of selection. We hypothesize that if linked selection reduced diversity at linked neutral sites along the genome, local levels of nucleotide diversity should be correlated with the rate of recombination and the density of targets of selection. In addition, we predict that if the efficiency of selection is a function of the demographic trajectory of populations during the Ice Age, large and more stable populations (i.e., larger long-term N_e_) will exhibit lower genetic load and a stronger correlation between nucleotide diversity and intrinsic genomic properties, such as recombination rate. These results have implications for our understanding of the relative importance of neutral and selective processes on the evolution of the genomic landscape of species heavily impacted by glaciations.

## Results

### Congruent population structure and genetic diversity

We characterized population genetic structure in Downy and Hairy Woodpeckers across an array of ecological zones that would have been subject to varying effects of Pleistocene climatic cycles. We collected whole genomes of 70 individuals each of Downy and Hairy Woodpecker (140 total samples; Table S1), representing seven geographic locations in North America: Northeast (NE), Southeast (SE), Midwest (MW), Southern Rockies (SR), Northern Rockies (NR), Pacific Northwest (NW), and Alaska (AK; Figure 1a–b). Sequenced reads were mapped to a pseudo-reference genome of Downy Woodpecker [52], yielding an average sequencing depth of 5.1x (1.4–12.5x) for Downy Woodpecker and 4.5x (1.1– 11.7x) for Hairy Woodpecker. A total of 16,736,465 and 15,463,356 single nucleotide polymorphisms (SNPs) were identified in the Downy and Hairy Woodpecker genomes, respectively, using the genotype likelihood approach implemented in ANGSD [53].

**Figure 1.**
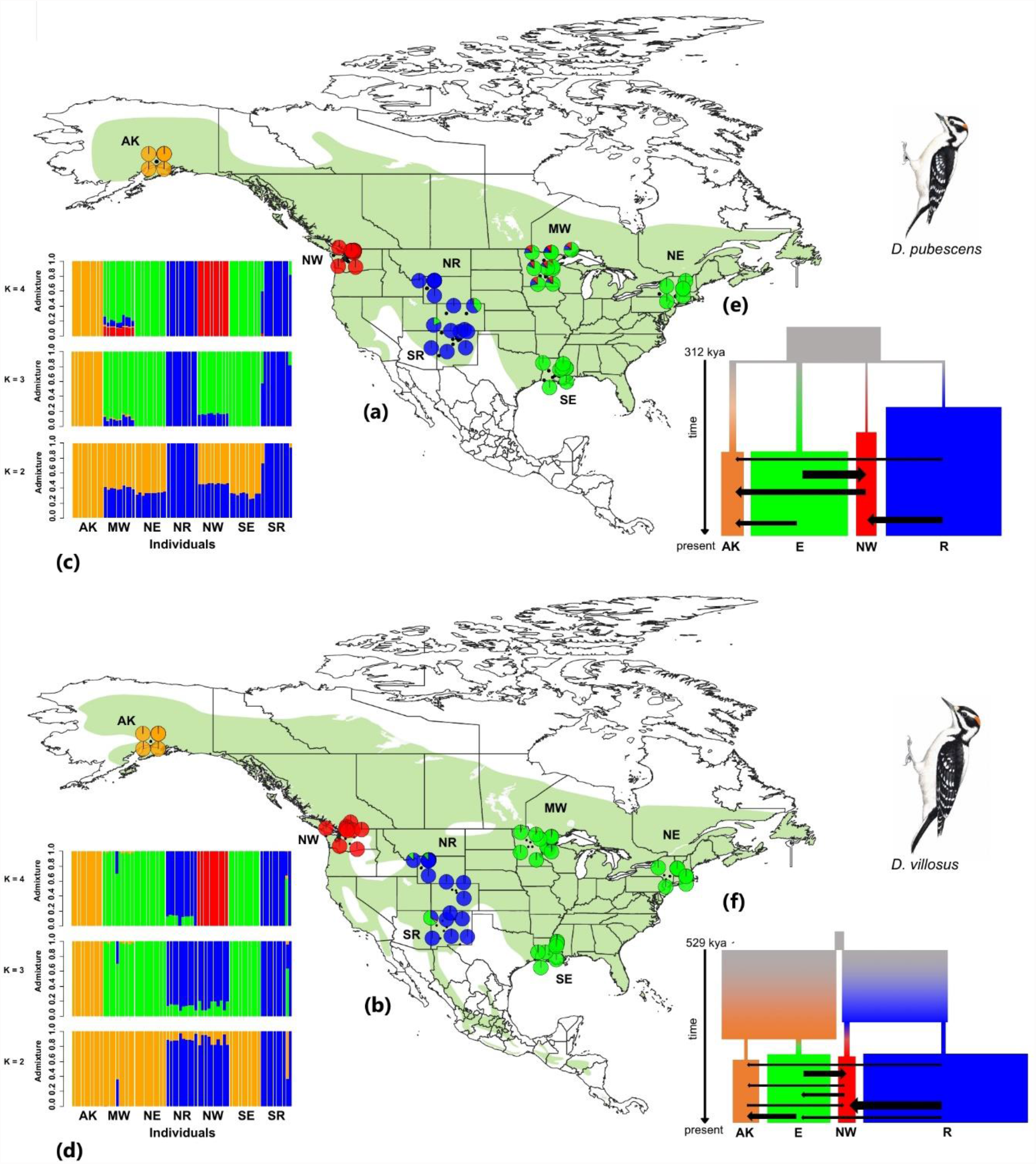
Geographic distribution of genetic variation and demographic history of the Downy (*D. pubescens*; top) and Hairy Woodpecker (*D. villosus*; bottom). **(a-b)** Results of the *NGSadmix* analysis for the K = 2–4. Each bar indicates an individual’s estimated ancestry proportion for each genetic cluster, represented by different colors. **(c-d)** Map indicating the current range of Downy and Hairy Woodpecker (green shade), the locality of the samples, and their respective admixture proportions from *NGSadmix* (pie charts). **(e-f)** The best-fit demographic models from *fastsimcoal2*. The width of the rectangles and arrows are scaled relative to the estimated effective population sizes in haploid individuals (N_e_) and the migration rate (*m*) in fraction of haploid individuals from donor population per generation, respectively. Only the values of migration rate > 10^−7^ ⨯ N_e_ migrants per generation are shown. Illustrations reproduced with permission from Lynx Edicions.

To assess patterns of genetic differentiation among these broadly distributed populations, we first performed a principal component analysis (PCA) on a subset of 71,229 and 71,816 independently-evolving (linkage disequilibrium r^2^ < 0.2) SNPs for Downy and Hairy Woodpecker, respectively. The three first principal components (PCs) explained together 7.5% (Downy) and 8.3% (Hairy) of the total genetic variance. We recovered congruent genetic structure across both species’ ranges (Figure 2a,c). Geographic structure was generally characterized by a genetic discontinuity between boreal-eastern and western populations. In Downy Woodpecker, however, the Pacific Northwest population fell more closely related to the Eastern group than the Western group (Figure 2a). Consistent with these findings, *NGSadmix* [54] supported four geographically congruent genetic clusters (K=4) in the Downy and Hairy Woodpecker: East (NE, SE, and MW), Pacific Northwest (NW), Rocky Mountains (SR and NR), and Alaska (AK; Figure 1c– d). The average genome-wide estimate of F_ST_ was slightly larger in Hairy Woodpecker (average F_ST_ = 0.1; 0.03–0.19) than Downy Woodpecker (average F_ST_ = 0.08; 0.03–0.16), indicating larger (but overlapping) levels of population differentiation. In both species, the largest values of F_ST_ involved comparisons between Alaska and other populations (Downy: F_ST_ [AK vs NR] = 0.16; Hairy: F_ST_ [AK vs SR] = 0.19), and the lowest were within the East and the Rocky Mountains clusters (Downy: F_ST_ = 0.03–0.06; Hairy: F_ST_ = 0.03– 0.04).

**Figure 2.**
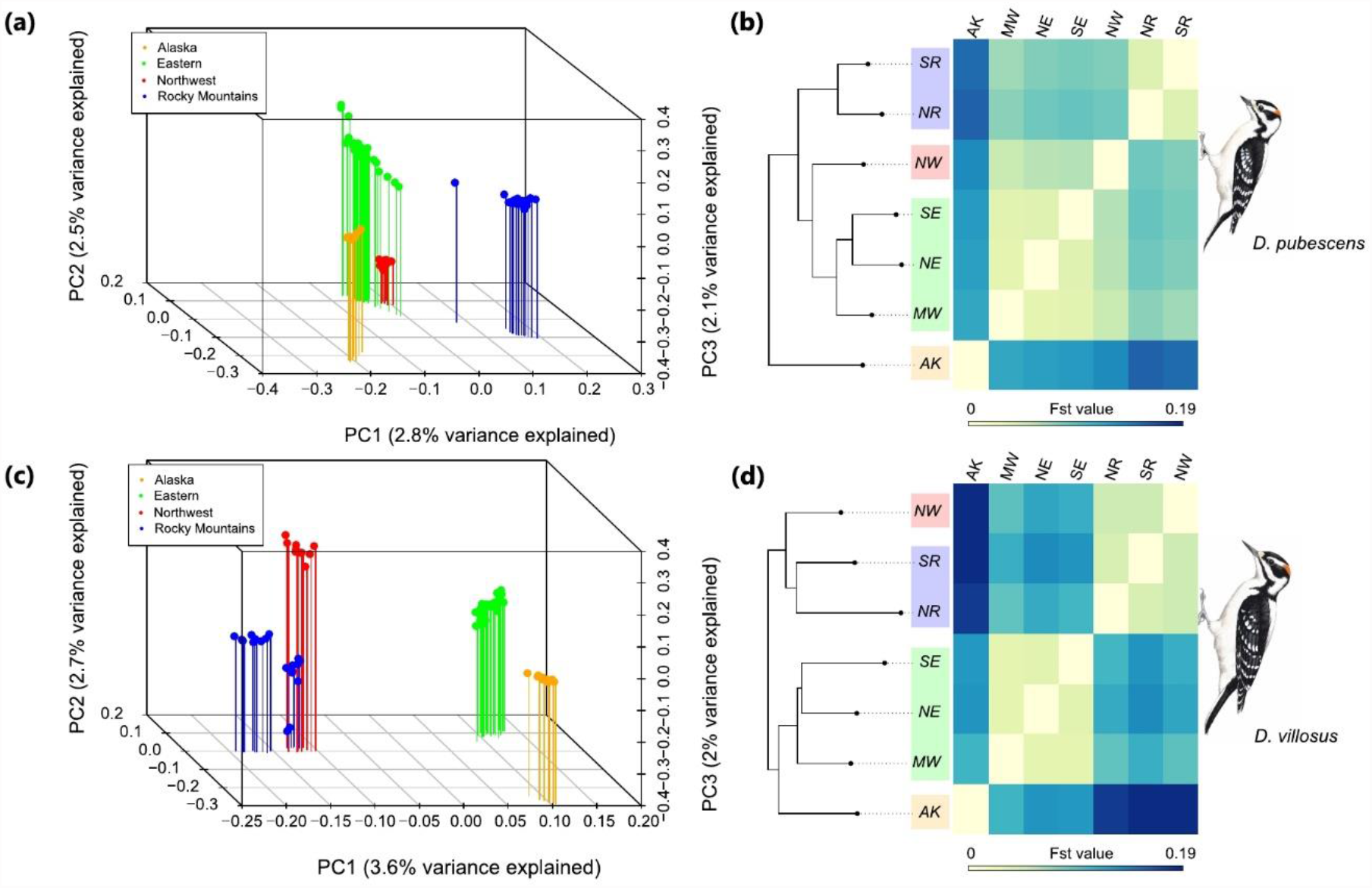
Population genetic structure in the Downy (top) and Hairy (bottom) Woodpecker. **(a**,**c)** Principal component analysis (PCA) of Downy and Hairy Woodpecker based on 71,228 and 71,763 unlinked genome-wide SNPs, respectively, with < 25% missing data and a minor allele frequency (maf) > 0.05. **(b**,**d)** Heatmap showing genome-wide pairwise F_ST_ values (left) and associated maximum likelihood tree based on the polymorphism-aware phylogenetic model (PoMo) in IQ-Tree 2. All nodes show 100% bootstrap support. Darker colors on the heatmap correspond to larger values of F_ST_. Illustrations reproduced with permission from Lynx Edicions.

Because the expansion and contraction of glaciers were expected to impact population structuring across the landscape, we explored spatial patterns of gene flow using the estimated effective migration surface (EEMS [55]). EEMS compares pairwise genetic dissimilarity among localities to identify geographic areas that deviate from the null expectation of isolation by distance (IBD). In both species, we detected a pronounced reduction in effective migration near the Great Plains and along the Rocky Mountains, especially in its Northern portion. In contrast, eastern North America showed a higher degree of connectivity when compared to the west (Figure 3). This finding indicates that major topographic features and variation in habitat availability contributed to the maintenance of population differentiation, despite the presence of gene flow.

**Figure 3.**
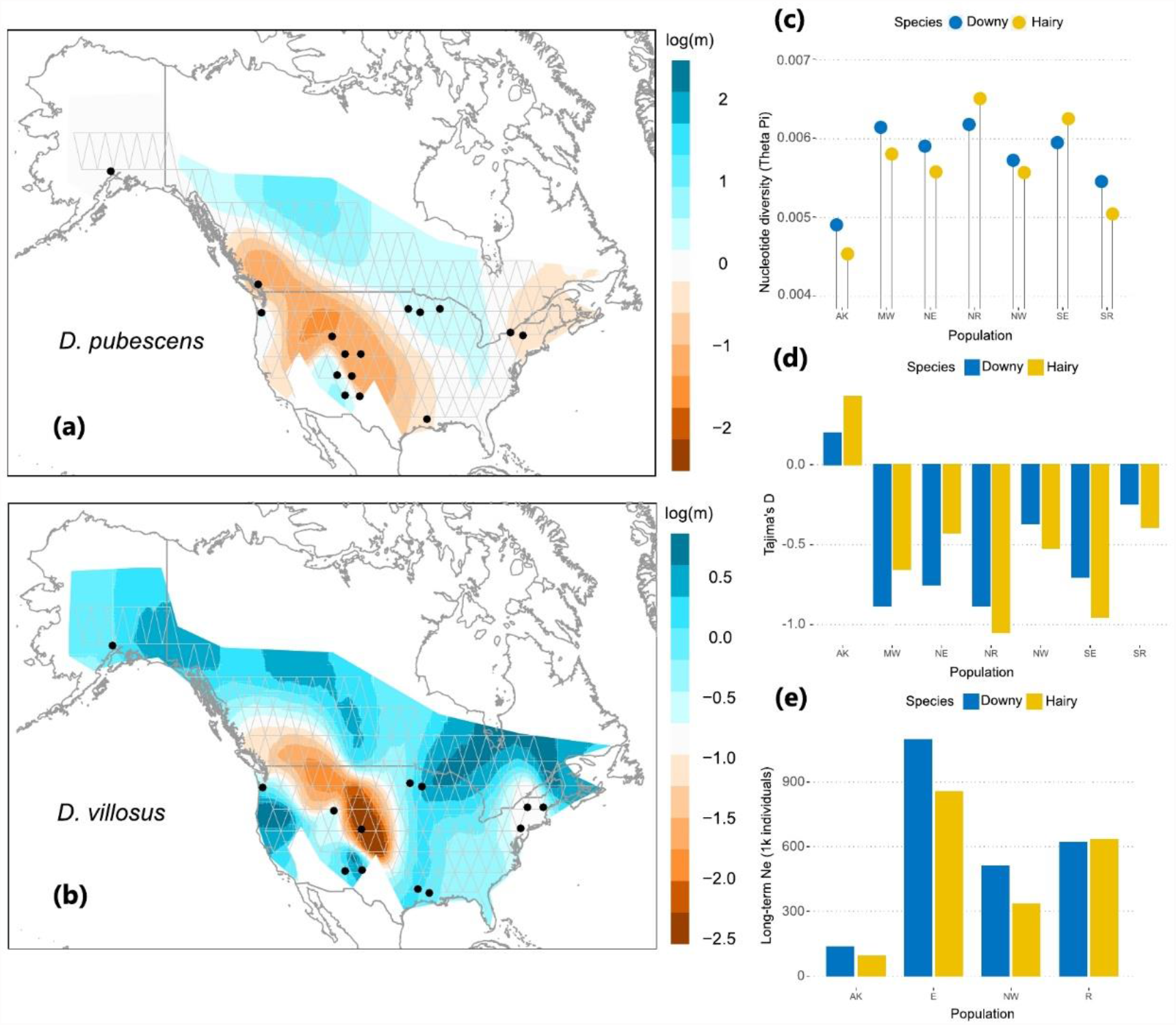
Spatial patterns of gene flow and genome-wide genetic variation in Downy and Hairy Woodpecker. **(a)** Effective migration surface inferred by EEMS in Downy Woodpecker and **(b)** Hairy Woodpecker. Warmer colors indicate lower and colder colors indicate higher effective migration (on a log scale) relative to the overall migration rate over the species range. Triangles represent the grid chosen to assign sampling locations to discrete demes. **(c)** Genome-wide pairwise nucleotide diversity (θ_π_) per population. **(d)** Genome-wide Tajima’s D per population. **(e)** The harmonic mean of effective population size (N_e_) estimated over the past one million years with Stairway Plot 2 for all four genetic clusters.

### Demographic history

We tested for signatures of Quaternary climatic oscillations on population dynamics of Downy and Hairy Woodpecker by assessing changes in N_e_ over time and estimating demographic parameters. First, we employed Stairway Plot 2 [56] to infer fluctuations in N_e_ over the past 500k years in each of the four detected genetic clusters assumed to represent panmictic populations. Stairway Plot 2 uses the site frequency spectrum (SFS) to fit a flexible multi-epoch model of changes in population size. For all demographic analyses, we used the folded SFS and specified a mutation rate of 4.007 × 10^−9^ mutations per site per generation and a generation time of one year for both species [57]. Changes in effective population size over time were generally consistent between both species, being characterized by recurrent episodes of bottleneck followed by population expansion (Figure 4a–b). We found that within each genetic cluster, nucleotide diversity was highly correlated with the harmonic mean of the N_e_ estimated from Stairway Plot 2 over the past 500 kya (long-term N_e_; linear regression: t = 4.876; R^2^ = 0.76; p < 0.002; Figure S1), indicating these independent analyses were consistent.

**Figure 4.**
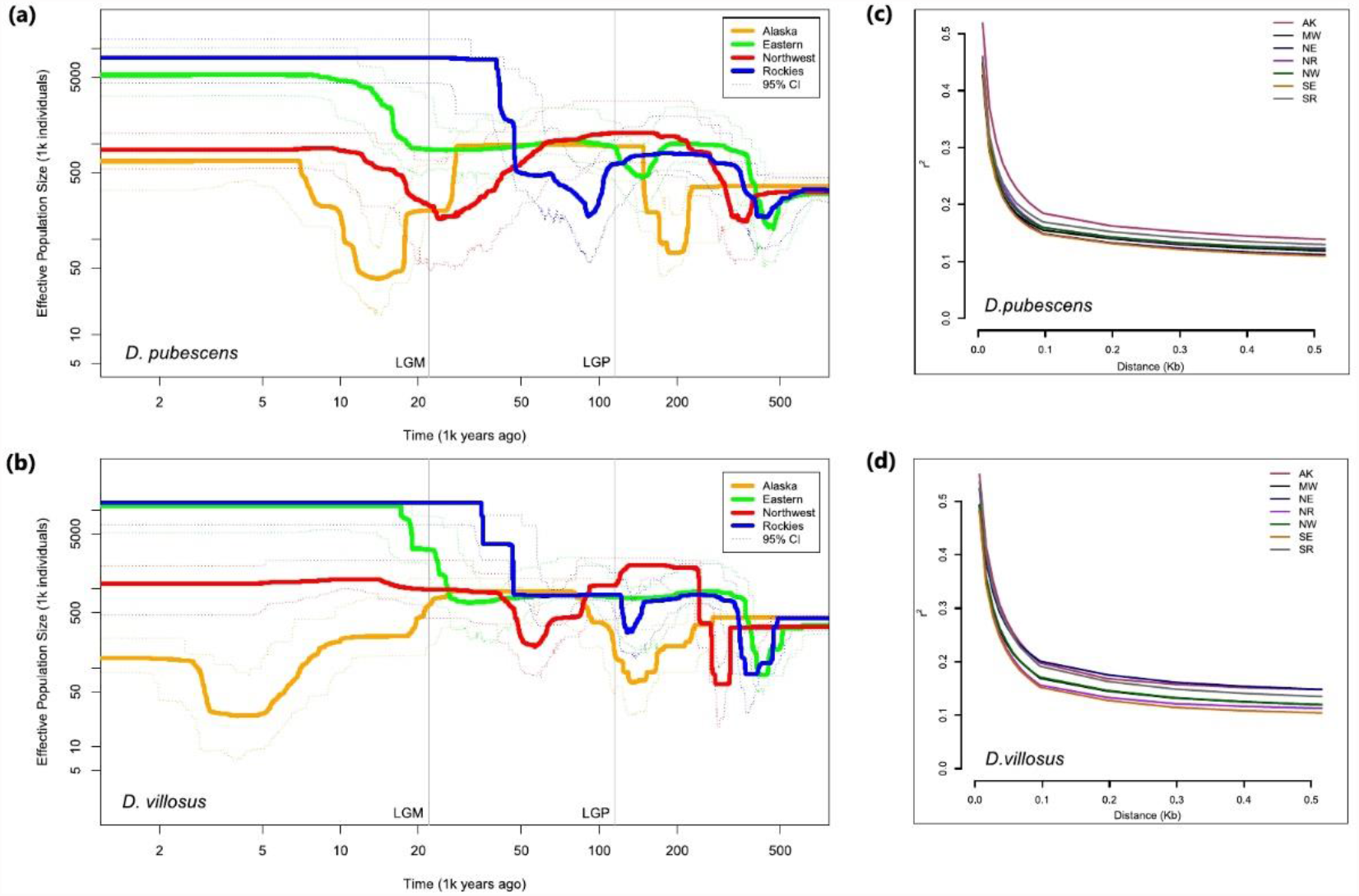
Changes in effective population size (N_e_) over time and linkage disequilibrium (LD) in Downy (top) and Hairy (bottom) Woodpecker. **(a–b)** Inferred history of effective population size of all four genetic clusters in Downy (a) and Hairy Woodpecker (b) obtained with Stairway Plot 2 using the folded SFS. For this analysis, we specified a mutation rate of 4.007 × 10^−9^ mutations per site per generation. Both axes are represented in a log scale. Dotted lines represent 95% confidence intervals, and vertical lines represent the Last Glacial Period (LGP; 115 kya) and the Last Glacial Maximum (LGM; 21 kya). **(c–d)** Decay of linkage disequilibrium (LD) in all seven populations of Downy (c) and Hairy (d) Woodpecker.

To further elucidate the evolutionary relationships among populations of Hairy and Downy Woodpecker, we built a rooted maximum likelihood tree from genome-wide intergenic SNPs using the IQ-TREE polymorphism-aware phylogenetic model (PoMo [58,59]). The topology for Hairy Woodpecker showed two distinct clades – an East + Alaska and a West clade. The tree for Downy Woodpecker, however, revealed a different topology. First, the Pacific Northwest population (NW) was more closely related to the eastern clade than to the western clade, supporting our PCA analysis. In addition, the Alaska (AK) population was sister to all other populations. Two hypotheses could explain this pattern: either (i) Alaska was a distinctive clade that differentiated from the other Downy Woodpecker populations as a consequence of persistence in a separate glacial refugium near Beringia, as has been suggested for other North American taxa [2,60,61], or (ii) the topology of the Downy Woodpecker population tree was more reflective of other factors, such as patterns of gene flow and geographic distance among localities, as opposed to the actual order of population splits. If this was the case, then we expect the relationships among populations to better fit a polytomous tree rather than a bifurcating tree.

To test these alternative population histories, we used the SFS-based method *fastsimcoal2* v2.6.0.3 [62] to estimate demographic parameters and evaluate the support for two alternative models – (i) a model where all populations diverge synchronously from a single ancestral refugium and expand independently with asymmetric gene flow, and (ii) a bifurcating model where populations diverge at different times from multiple refugia (e.g., Beringia and East or East and West) and expand independently with asymmetric gene flow, following the IQ-TREE topology.

Demographic analyses with *fastsimcoal2* show differing support for alternative demographic models among the two focal species. The best-supported model for Hairy Woodpecker was model ii (Table S2; Figure 1f), in which two ancestral populations (putatively located in East and West) diverged from each other around 529 kya (95% CI = 513–561 kya; Table S3) and gave rise to the four genetic clusters, which then underwent strong bottlenecks. A final explosive expansion then occurred between 193–212 kya when populations grew up to 12-fold. In contrast, Downy Woodpecker showed support for model i, in which all populations diverge from a single major refugium (Table S2; Figure 1e). This divergence occurred around 312 kya (95% CI = 146–551 kya; Table S3) and was accompanied by a large bottleneck, reducing N_e_ to less than 10% of its original size in most populations. A final population expansion then occurred at the end of the Mid-Pleistocene (152–232 kya). Overall, estimates of N_e_ from *fastsimcoal2* confirmed the trends observed in Stairway Plot 2, albeit with less resolution. We found large and variable levels of post-expansion gene flow across populations in both species (Downy: 0–4.8 migrants per generation; Hairy: 0– 6.66 migrants per generation) that confirmed our EEMS migration surfaces.

### Genomic correlates of nucleotide diversity and differentiation

To elucidate the evolutionary processes shaping levels of genetic variation along the genome of Downy and Hairy Woodpecker, we investigated the correlation between regional levels of nucleotide diversity, measured across non-overlapping 100 kb windows, and three genomic features: recombination rate, gene density, and base composition. We found that nucleotide diversity varied widely along the genome (θ_π Downy_ = 7.5 ⨯ 10^−4^–1.9 ⨯ 10^−2^; θ_π Hairy_ = 1.1 ⨯ 10^−3^–2.2 ⨯ 10^−2^), but this variation was highly correlated between Downy and Hairy Woodpecker (Pearson’s *r* = 0.9; p < 0.001; Figure S2). To estimate recombination rates, we used ReLERNN [63], a method that uses a machine-learning approach to infer per-base recombination rates. We found recombination rates to be highly correlated between the two species (Pearson’s *r* = 0.66; p < 0.001). Across the genome, we estimated a mean per-base recombination rate (*r*) = 2.42 × 10^−9^ c/bp (0– 3.87 ⨯ 10^−9^) in Downy Woodpecker and *r* = 3.69 × 10^−9^ c/bp (4.85⨯10^−10^–3.87 ⨯10^−9^) in Hairy Woodpecker. Considering the average long-term N_e_ of Downy and Hairy Woodpecker as approximately 1 × 10^6^ in the East population, these recombination rates correspond to a population-scaled rate ρ = 4N_e_r = 0.008 and 0.012, respectively. Mean recombination rates were 2–3-fold higher in autosomal chromosomes compared to the sex-linked Z chromosome (Figure S3–4), consistent with differences in N_e_ between sex chromosomes [64–66]. As a result of both high recombination rates and large N_e_, we also observed that linkage disequilibrium (LD) in Downy and Hairy Woodpecker decays very rapidly. LD drops to half of its initial levels in less than 100 bp (Figure 5c–d). Consistently, the average LD was greater for populations with smaller N_e_ or populations that have likely experienced a more recent founder event, such as Alaska and the Southern Rockies. We found a significant positive association between nucleotide diversity (θ_π_) and recombination rates in both species (linear regression – Downy: t = 47.67, R^2^ = 0.165, p < 0.001; Hairy: t = 54.17, R^2^ = 0.204, p < 0.001; LOESS regression – Downy: span = 0.5, R^2^ = 0.207; Hairy: span = 0.5, R^2^ = 0.207; Figure 5a–b). This association, however, is expected (to a certain extent) even if diversity is not correlated with recombination rates because recombination rates are estimated directly from θ_w_.

**Figure 5.**
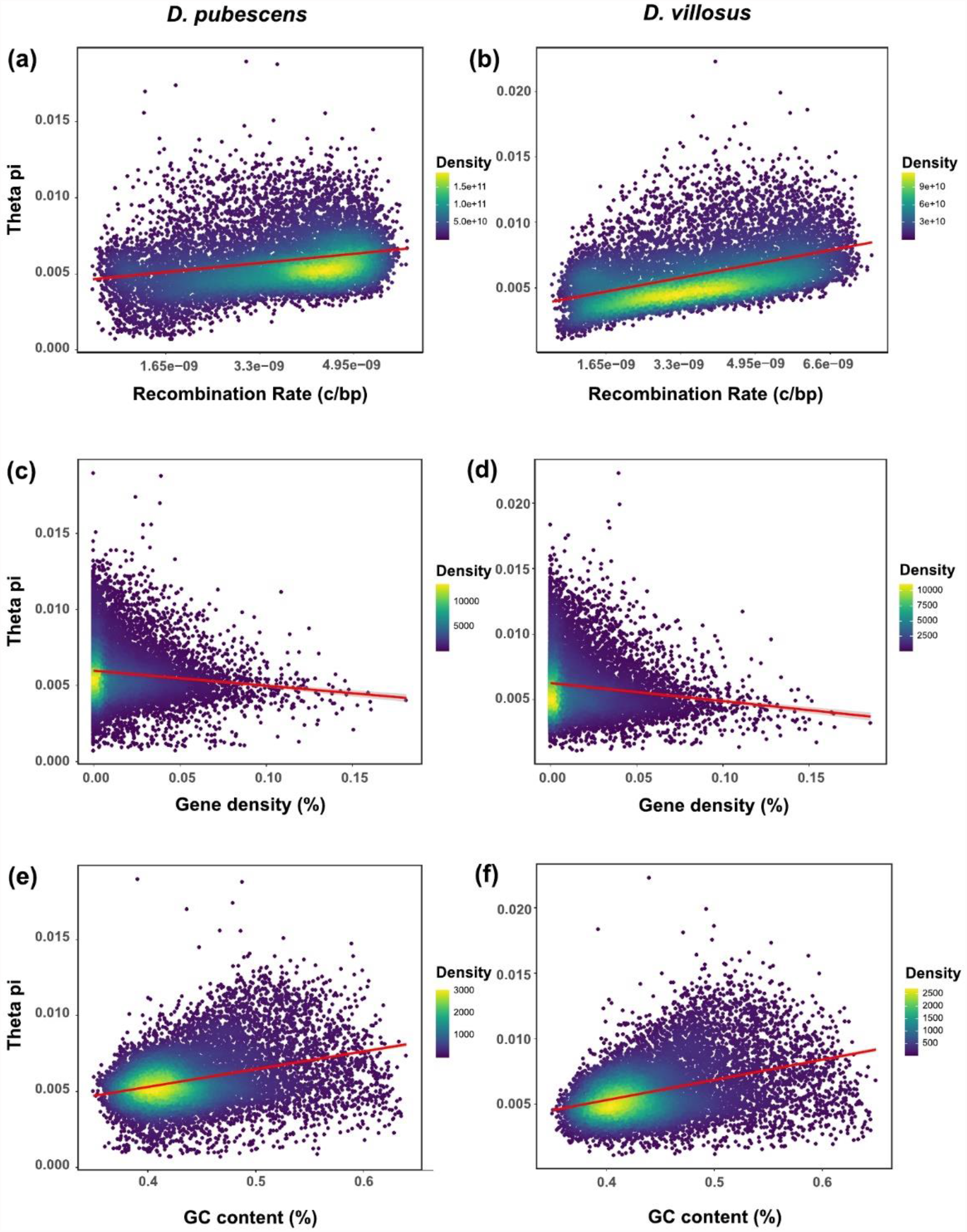
Genomic predictors of nucleotide diversity in Downy (left) and Hairy (right) Woodpecker. Association between nucleotide diversity (θ_π_) and three features of the genome: **(a–b)** recombination rates (Downy: t = 47.67, p <0.001; Hairy: t = 54.17, p <0.001), **(c–d)** gene density (Downy: t = -12.03, p <0.001; Hairy: t = -14.89, p <0.001), and **(e–f)** GC content (Downy: t = 36.37, p <0.001; Hairy: t = 44.16, p <0.001). Each point in the scatter plot represents a 100 kb window of the genome. Colors indicate the density of points.

To further investigate the impact of linked selection on the genomic landscape of diversity, we also tested the prediction that regions of the genome with a higher density of targets of selection (i.e., genes) exhibit lower nucleotide diversity. Gene density was measured as the percentage of coding sequence in each of the 100 kb windows. Our results revealed a weak but significant negative association between nucleotide diversity (θ_π_) and gene density (linear regression – Downy: t = -12.03, R^2^ = 0.0123, p < 0.001; Hairy: t = -14.89, R^2^ = 0.0189, p < 0.001; LOESS regression – Downy: span = 0.5, R^2^ = 0.0139; Hairy: span = 0.5, R^2^ = 0.021; Figure 5c–d). This association was not driven by the collinearity between gene density and recombination because this correlation was positive and negligible (Downy: Pearson’s *r* = 0.045, p < 0.001; Hairy: Pearson’s *r* = 0.032, p < 0.001). We also found that regions with high GC content tended to show higher nucleotide diversity (linear regression – Downy: t = 36.37, R^2^ = 0.0123, p < 0.001; Hairy: t = 44.16, R^2^ = 0.145, p < 0.001; Figure 5e–f). GC content, however, was positively correlated with gene density in both species (Downy: Pearson’s *r* = 0.25; p < 0.001; Hairy: Pearson’s *r* = 0.25; p < 0.001; Figure S5–6) and weakly correlated with recombination rates in Hairy Woodpecker (Pearson’s *r* = 0.064; p < 0.001; Figure S5–6). We then performed a principal component regression (PCR) to separate the effect of individual explanatory variables and control for the multicollinearity among predictor variables. Principal component regression summarizes variables into orthogonal components (PCs) and uses these components as predictors in a linear regression. PC2, which represented almost exclusively recombination rates (Table 1), uniquely explained 12.3% and 18.6% of variation in nucleotide diversity in Downy and Hairy Woodpecker, respectively (PC2 linear regression – Hairy: t = 51.1, R^2^ = 0.186, p < 0.001; Downy: t = 40.14, R^2^ = 0.123, p < 0.001). Both PC1 and PC3 represented the correlation between gene density and GC content, but PC3 had a much stronger effect (Table 1), accounting for 14.4% and 15.5% of the variation in nucleotide diversity in Downy and Hairy Woodpecker, respectively (PC3 linear regression – Downy: t = 45.92, R^2^ = 0.155, p < 0.001; Hairy: t = 43.97, R^2^ = 0.144, p < 0.001). Considering that gene density and GC content had an equal contribution to PC3 (Table 1), we were unable to differentiate their relative contributions to the relationship. Regardless, our analyses confirm the central role that these genomic properties played in shaping patterns of nucleotide diversity along the genome.

**Table 1.** Principal component regression.

| Species | Explanatory variables | % of variance explained ($R^2$) | | |
| --- | --- | --- | --- | --- |
|  |  | PC1 | PC2 | PC3 |
| Downy Woodpecker | Recombination rate | 0.08 | 11.89 | 0.11 |
|  | Gene density | 1.7 | 0.03 | 7.8 |
|  | GC content | 1.65 | 0.36 | 7.58 |
|  | Total | 3.45 | 12.3 | 15.51 |
| Hairy Woodpecker | Recombination rate | 0.35 | 17.35 | 0.01 |
|  | Gene density | 2.84 | 1.02 | 7 |
|  | GC content | 2.96 | 0.06 | 7.34 |
|  | Total | 6.18 | 18.6 | 14.47 |

The effect of linked selection is expected to be weaker in populations that underwent more severe bottlenecks due to their smaller long-term N_e_ when compared to stable populations that maintained large N_e_ [67,68]. We tested this prediction by quantifying the strength of correlation between nucleotide diversity (θ_π_) and gene density in all four genetic clusters of Downy and Hairy Woodpecker which showed varied demographic responses to the Pleistocene glaciations. We found that long-term N_e_ predicted the coefficient of correlation between genetic diversity and the density of targets of selection (Table 2). Alaska, for example, showed the weakest correlation (Downy: Pearson’s *r* = -0.1008, t = -10.8, p < 0.001; Hairy: Pearson’s *r* = -0.1083, t = -11.6, p < 0.001), whereas Rocky Mountains showed the strongest (Downy: Pearson’s *r* = -0.1106, t = -11.9, p < 0.001; Hairy: Pearson’s *r* = -0.1351, t = -14.5, p < 0.001). Although the differences in coefficients were small, these results support the expectation that different demographic trajectories affect the efficacy of natural selection owing to differences in levels of genetic drift.

**Table 2.** Strength of correlation between nucleotide diversity (θ_π_) and gene density across the four genetic clusters of Downy and Hairy Woodpecker.

| Populations | Downy Woodpecker |  | Hairy Woodpecker |  |
| --- | --- | --- | --- | --- |
| | Pearson's $r$ | t-value | Pearson's $r$ | t-value |
| AK | -0.1008* | -10.867 | -0.1083* | -11.643 |
| NW | -0.1007* | -10.847 | -0.1384* | -12.966 |
| E | -0.1077* | -11.618 | -0.1215* | -13.084 |
| R | -0.1106* | -11.927 | -0.1351* | -14.571 |

\* $p < 0.001$ .

Because genomic properties are also expected to impact levels of population differentiation across the genome, we also tested the association between nucleotide diversity, recombination rate, and the average intraspecific population differentiation (F_ST_) across non-overlapping 100 kb windows. For each window, we calculated the F_ST_ between each pair of populations and summarized the global F_ST_ landscape using two approaches: (i) the average F_ST_ across all population pairs; and (ii) the first principal component (PC1) explaining most of the variation in pairwise F_ST_ (Downy: variance explained = 37.51%; Hairy: variance explained = 47.5%). Summaries of F_ST_ produced by these two approaches were highly correlated (Downy: Pearson’s *r* = 0.97; p < 0.001; Hairy: Pearson’s *r* = 0.98; p < 0.001), so we only considered the average F_ST_ for simplicity. There was considerable variation in F_ST_ along the genome (Downy: F_ST_ = 0.01– 0.25; Hairy: F_ST_ = 0.01–0.32), indicating high variability in patterns of population differentiation. We recovered a significant negative association between average F_ST_ and nucleotide diversity, suggesting that areas of genome that show elevated differentiation tend to be characterized by reduced diversity (linear regression – Downy: t = -19.12, R^2^ = 0.03; p < 0.001; Hairy: t = -53.49, R^2^ = 0.2; p < 0.001; Figure 6). Finally, we found a negative association between average F_ST_ and recombination rates, indicating higher differentiation in regions of low recombination (linear regression – Downy: t = -32.18, R^2^ = 0.08; p < 0.001; Hairy: t = -41.55, R^2^ = 0.13; p < 0.001).

**Figure 6.**
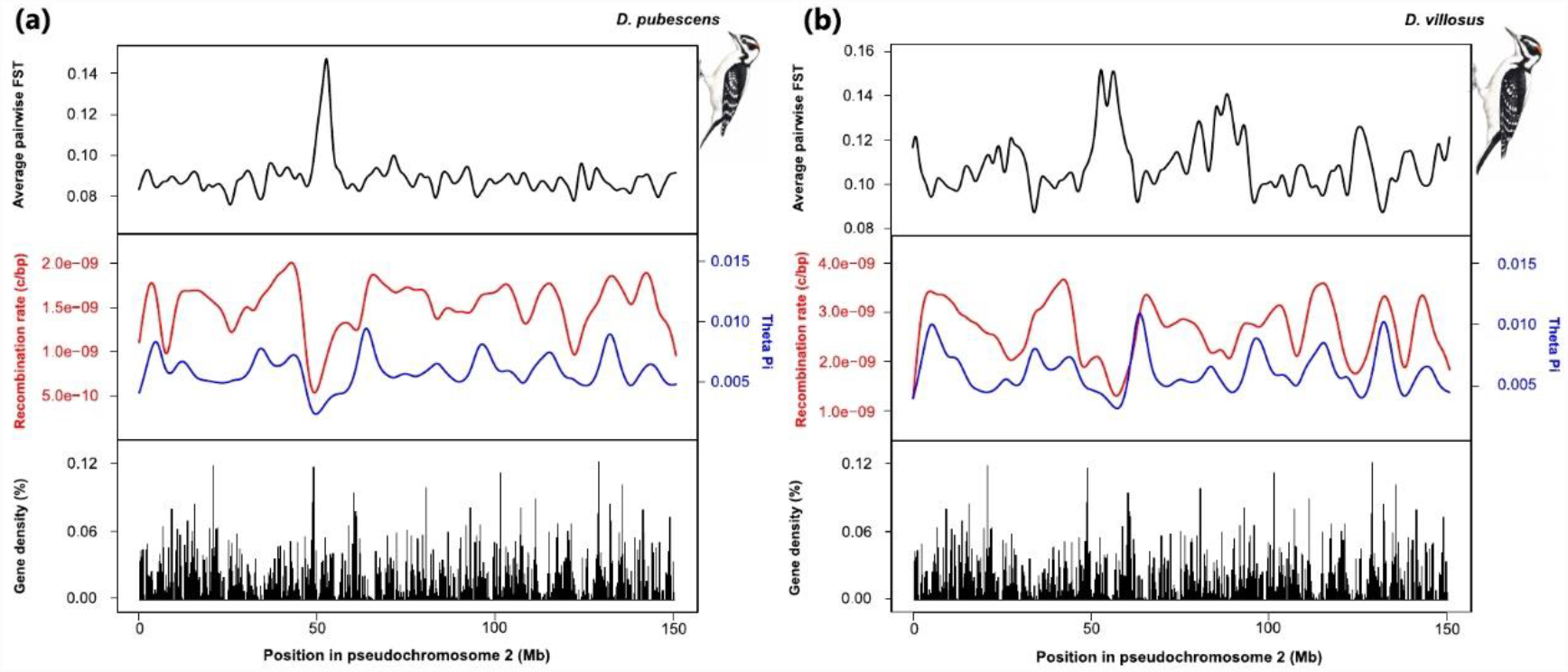
Landscape of diversity and differentiation of chromosome 2 of Downy (a) and Hairy (b) Woodpecker for illustration. Top plot shows the average pairwise F_ST_ calculated across non-overlapping 100 kb windows. Middle plot indicates the recombination rate in c/bp (red) and the nucleotide diversity (θ_π_; blue) for each non-overlapping 100 kb window. Bottom plot represents the percentage of coding sequence in each non-overlapping 100 kb window. Illustrations reproduced with permission from Lynx Edicions.

### Genetic load and the efficacy of selection

To further explore the magnitude of linked selection in the genome of Downy and Hairy Woodpecker, we classified each variant according to their functional impact as predicted by the gene annotation. We found that the majority of identified SNPs in Downy and Hairy Woodpecker were classified as modifiers (Downy: 99.35%; Hairy: 99.13%), which are variants in intergenic or intronic regions whose impacts are hard to determine but tend to be neutral to nearly neutral. Low impact variants (i.e., synonymous mutations) characterized 0.46% and 0.64% of SNPs in Downy and Hairy Woodpecker, respectively. Moderate impact variants, mutations that cause a change in amino acid sequence (i.e., nonsynonymous mutations) represented 0.17% and 0.22% of the SNPs in Downy and Hairy Woodpecker, respectively. Finally, only 0.006% and 0.007% of the SNPs were classified as high impact in Downy and Hairy Woodpecker, respectively. These variants correspond to mutations that cause loss of function, such as loss or gain of a start or stop codon and are therefore expected to occur at very low frequencies.

We investigated differences in the burden of deleterious alleles carried by populations of Downy and Hairy Woodpecker that could reflect differences in the efficacy of purifying selection. For this analysis, we focused on sites that were polymorphic in at least one of the two species and whose ancestral states could be determined unambiguously. Our results revealed that the frequency distribution of mutations with moderate and high impact shifted downwards compared to the mutations with low impact (Figure 7a–b). This indicates that purifying selection was successful in purging mutations that were highly deleterious. Hairy Woodpecker, however, showed a larger excess of low frequency mutations of high impact when compared to Downy Woodpecker (Figure 7a–b), suggesting that purifying selection might have been more efficient in Hairy Woodpecker. To further evaluate whether the efficacy of purifying selection varied across populations with different demographic trajectories, we estimated for each individual the genetic load as the ratio of the count of homozygous derived alleles of high impact (i.e., highly deleterious) over the count of homozygous derived alleles of low impact (i.e., synonymous). This metric is a proxy for the genetic load under a recessive model while controlling for the underlying population differences in the neutral SFS [69,70]. We also computed the same metric considering an additive model, in which the presence of a single copy of the derived allele has fitness consequences. Our results reveal that the recessive deleterious load was overall larger in Downy than Hairy Woodpecker, but this difference was not statistically significant (Kruskal-Wallis χ^2^ = 1.33, df =1, p = 0.24; Figure 7c–d). The recessive deleterious load was much larger in the Rocky Mountains when compared to other populations. Alaska also showed elevated recessive deleterious load in both species, generally larger than the East and Pacific Northwest (Figure 7c–d). Overall, these findings do not support the prediction that populations with stronger bottlenecks exhibit high deleterious load.

**Figure 7.**
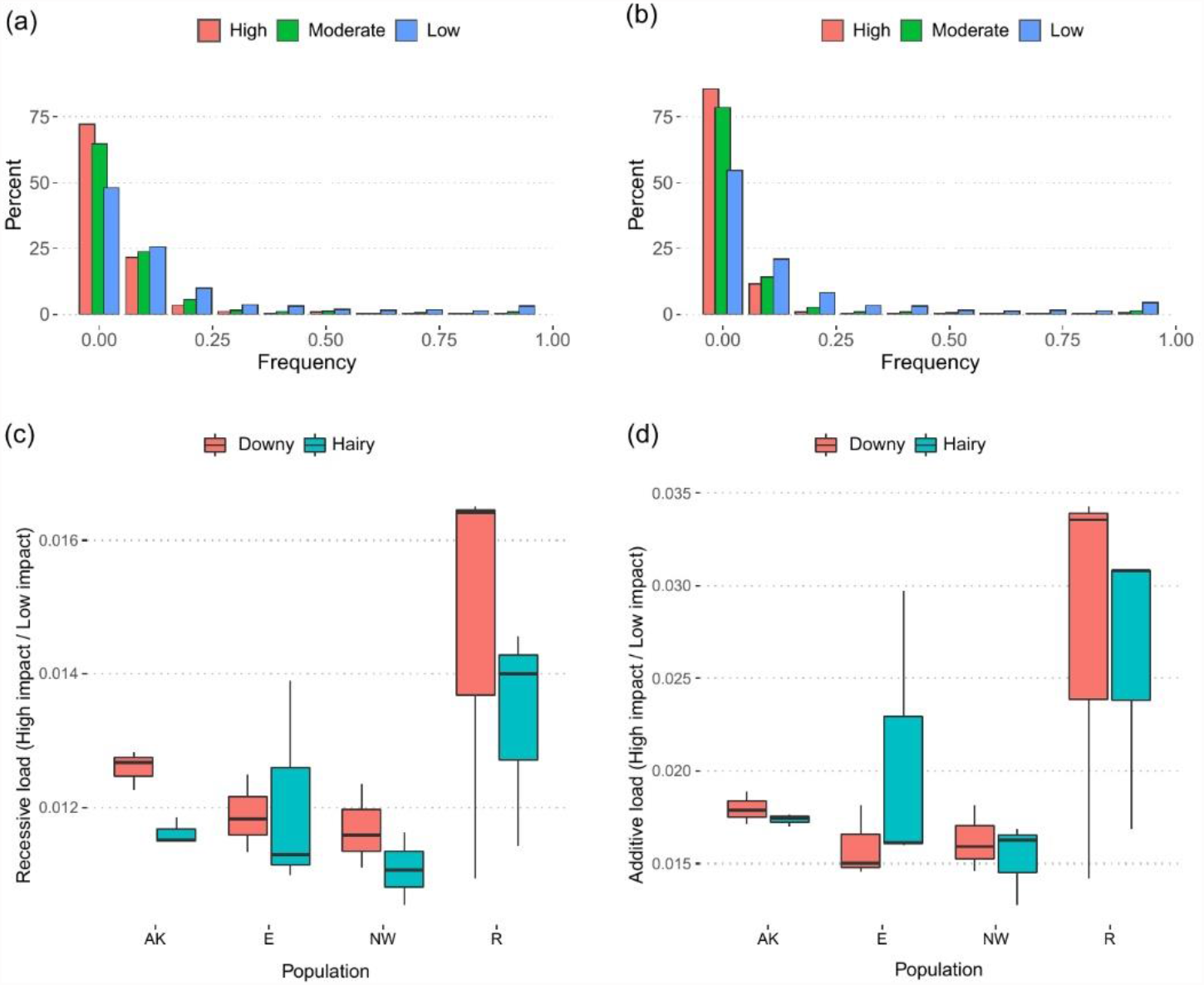
Deleterious load in Downy and Hairy Woodpecker. **(a)** Site frequency spectrum (SFS) for variants with low (neutral), moderate (mild), and high (deleterious) impact in Downy Woodpecker and **(b)** Hairy Woodpecker. **(c)** Ratio of homozygous derived variants of high impact (deleterious) over homozygous derived variants of low impact (neutral) in each genetic cluster and species (recessive model). **(d)** Ratio of the total number of derived variants of high impact (deleterious) over total number of derived variants of low impact (neutral) in each genetic cluster and species (additive model). Horizontal bars denote population medians.

Lastly, we investigated the overall impact of natural selection on protein-coding sequences of Downy and Hairy Woodpecker. We calculated the ratio of synonymous over nonsynonymous substitutions (dN/dS) along the branches leading to Downy and Hairy Woodpecker using a set of 397 high-quality orthologous genes distributed throughout the genome. dN/dS ratio was higher in Downy Woodpecker (dN/dS = 0.065) than in Hairy Woodpecker (dN/dS = 0.053), suggesting that purifying selection might have been weaker in the Downy Woodpecker lineage over deeper evolutionary times (i.e. >4N_e_ generations ago [71–73]).

## Discussion

Our genomic analyses reveal that both Ice Age demographic fluctuations and linked selection played a significant role shaping patterns of diversity and differentiation across populations and along the genomes of Downy and Hairy Woodpecker. We found that genome-wide nucleotide diversity, as well as the landscape of recombination, are highly correlated between these two species, which diverged more than 8 mya. Despite shared environmental pressures, this coupling suggests that intrinsic properties of the genome, such as recombination rate, might be conserved across deep evolutionary time. We posit that linked selection might underlie the genomic heterogeneity observed, as demonstrated by a significant association between nucleotide diversity, recombination rate, and gene density. Despite strong fluctuations in N_e_ over the Pleistocene, Downy and Hairy Woodpecker maintained large population sizes, which might have facilitated the action of natural selection. Nevertheless, given the large differences in long-term N_e_ observed among populations, our results indicate variation in the efficacy of selection.

### Conserved properties of the genome underlie the correlated genomic landscape of Hairy and Downy Woodpecker

We recovered large heterogeneity in patterns of nucleotide diversity (θ_π_) and F_ST_ along the genomes of Downy and Hairy Woodpecker. Despite this variation, our results revealed a highly correlated genomic landscape between the two species. Such covariation in levels of genome-wide measures of diversity and differentiation across distantly related species is common [34,35,74–77] and suggests that properties of the genome, such as mutation rate, recombination rate, and density of targets of selection are conserved across deep evolutionary time [78]. For example, bird genomes are known to show large karyotypic stability, with very few chromosomal rearrangements and high synteny across highly divergent species [79–82]. Features of the genome, such as recombination rates and GC content, might also be conserved across species. We found that estimates of recombination rate are highly correlated between Downy and Hairy Woodpecker, although higher in the latter. Linkage disequilibrium (LD), which is a function of both recombination rate and N_e_, was extremely short in Downy and Hairy Woodpecker. Whereas linkage disequilibrium extends for over thousands of base pairs in humans [83,84], for instance, it breaks after only 100 bp in Downy and Hairy Woodpecker. Such properties have been observed in other bird species with very large N_e_ [85,86]. We also found large variation in recombination rates both within and among chromosomes, with the Z chromosome showing the lowest rates. Considering the lack of recombination across much of the Z chromosome in female birds (heterogametic sex; ZW), at the population level, crossing-over occurs at a much lower rate in sex chromosomes than in their autosome counterparts [64,87,88]. Similar to Downy and Hairy Woodpecker, recombination in the chicken (*Gallus gallus*) was approximately 2.5 times lower in the Z chromosome than in the autosomes [89,90]. As a consequence, many bird species show reduced diversity and faster divergence in the Z chromosome [64,85,91,92].

### The interplay between natural selection and recombination produces a heterogeneous genomic landscape

One of the main mechanisms proposed to explain the substantial heterogeneity in levels of polymorphism along the genome is the effect of linked selection [21,23,24]. Both positive selection (i.e., in favor of a beneficial allele) and negative selection (i.e., against a deleterious allele) are expected to reduce diversity around functional elements [21,23]. Such a reduction is extended to all neighboring sites that happen to be linked to the target of selection (hitchhiking effect [21]). The extent to which adjacent sites are affected by linked selection is dependent on the recombination landscape, such that regions where recombination rate is lower tend to show lower genetic diversity and vice versa [32,93,94]. Similarly, the higher the density of functional elements (i.e., targets of selection), the more severe is the reduction in genetic diversity due to the effect of recurrent selection [33,38–40]. Although a correlation between nucleotide diversity and recombination may arise in the absence of linked selection, we do not expect that to be true for gene density, since directional selection unavoidably affects levels of polymorphism in regions presumed to be functional. Such correlations are therefore interpreted as evidence of the effect of selection on linked neutral sites and can be used to assess the magnitude of linked selection [24,95]. In light of these results, we identified strong evidence that linked selection has contributed to patterns of genetic diversity along the genomes of Downy and Hairy Woodpecker. First, nucleotide diversity (θ_π_) was positively associated with recombination rates in both species. Second, there was a weak but highly significant association between nucleotide diversity (θ_π_) and gene density. Third, as predicted by theory, the strength of association between nucleotide diversity (θ_π_) and gene density varied according to the long-term N_e_, such that larger populations showed more pronounced signatures of linked selection.

Natural selection is also expected to impact levels of genetic differentiation along the genome [35,96,97]. We estimated a weak but significant negative association between nucleotide diversity (θ_π_) and the average pairwise F_ST_, indicating that regions of the genome that are highly differentiated between populations tend to show reduced diversity. These correlations are consistent with the effect of linked selection continuously eroding diversity near targets of selection (especially in regions of low recombination), which leads to the inflation of local levels of population differentiation [96]. Because beneficial alleles are not expected to appear frequently, background selection against deleterious alleles is the most likely selective mechanism underlying the correlation between F_ST_, nucleotide diversity, and recombination rate [97,98]. These findings suggest that population-specific selection associated with local adaptation (i.e., divergent selection) is not necessary to produce a correlated genomic landscape. Comparative analyses across both distantly and closely related bird species demonstrate that linked selection can reduce genetic diversity prior to population splits and consequently produce parallel patterns of genetic differentiation in regions of low recombination [75,77,98,99].

### Dynamic population demography characterizes the evolution of Hairy and Downy Woodpecker in the Pleistocene

We found that population structure was spatially congruent between Downy and Hairy Woodpecker, but that their demographic histories and extent of genetic structuring varied. Both species were characterized by phylogeographic clusters that are consistent with previous studies [12,48,49] and were concordant with structuring in glacial refugia, albeit forming during different time scales. The observation of common geographic patterns formed across different time periods highlights the predictability of the interaction of the physical landscape, drift, and gene flow on genetic diversity. Further evidence of the dramatic effects of Pleistocene climatic fluctuations on genetic diversity were the repeated cycles of population contraction and expansion. Yet, despite strong variation in N_e_ over the past 500 ky, our data indicates that Downy and Hairy Woodpecker have been resilient enough to maintain relatively large populations, which favored the maintenance of very high genetic diversity, even in the face of repeated bottlenecks. While non-equilibrium population dynamics are a hallmark of species that occur in previously glaciated areas [2], the relationship between the magnitude of Plesitocene population size reductions and the efficacy of selection under these conditions remain poorly explored in empirical systems.

Consistent with theoretical predictions, nucleotide diversity within populations was strongly correlated with the long-term N_e_. Alaska showed the lowest genome-wide genetic diversity, likely as a consequence of being one of the latest areas to be deglaciated and most recently founded. On the other hand, the Northern Rockies exhibited the largest nucleotide diversity and long-term N_e_, in both focal species. Data from multiple sources support the existence of a temporally fluctuating ice-free corridor along the Canadian Rocky Mountains that might have functioned as a glacial refugium [10,100–102]. Thus, it is possible that suitable habitat might have allowed rapid growth and persistence of large populations in the North Rockies during the glacial periods of the Pleistocene.

### The efficacy of linked selection was affected by different evolutionary trajectories of Downy and Hairy Woodpecker

We investigated whether differences in the demographic trajectories of populations of Downy and Hairy Woodpecker in response to the Pleistocene glaciation had an impact on the efficacy of natural selection across the genome. Given that purifying selection is more efficient in larger populations [103], we hypothesized that populations that underwent a stronger bottleneck or maintained lower levels of N_e_ were more likely to have accumulated highly deleterious mutations (i.e., genetic load [26–30]). We failed to find support for this prediction. In contrast to our expectations, we found that the Rocky Mountains, the genetic cluster with the largest long-term N_e_, exhibited the largest genetic load in both species. One possible explanation for this finding is that highly deleterious alleles might have been more efficiently purged from populations that went through more severe bottlenecks due to higher inbreeding [67]. For example, species whose populations underwent extreme bottlenecks show fewer mutations of high impact because extensive inbreeding makes highly deleterious alleles more likely to be exposed in homozygosity [104–106]. This is not the case for Downy and Hairy Woodpecker, which despite repeated episodes of bottlenecks still managed to maintain considerably large population sizes, making inbreeding very unlikely to have occurred. Besides, we found that Alaska, the population with the lowest long-term N_e_, does not carry the fewest highly deleterious alleles, as predicted by the” purging under inbreeding” scenario. Instead, it carries a larger load than the East and the Pacific Northwest, which are populations with a higher long-term N_e_. At the species level, however, we found that genetic load was generally larger in Downy Woodpecker than Hairy Woodpecker, which is consistent with more efficient purifying selection in Hairy Woodpecker. This finding makes sense considering that Hairy Woodpecker exhibits slightly larger N_e_ than Downy Woodpecker. Supporting this observation, we also found a larger excess of highly deleterious mutations at low frequencies in Hairy Woodpecker, indicating that deleterious alleles were less likely to rise to high frequencies in Hairy Woodpecker than Downy Woodpecker likely due to more efficient selection. Lastly, we observed that the genome-wide ratio of non-synonymous over synonymous substitutions (dN/dS) was higher in Downy Woodpecker than Hairy Woodpecker. Elevated genome-wide, as opposed to gene-specific, dN/dS ratio is suggestive of a reduction in the efficacy of purifying selection [71,72]. This result indicates that a smaller N_e_ in the lineage leading to Downy Woodpecker might have allowed more fixation of slightly deleterious alleles.

In conclusion, we investigated the impact of demography and natural selection on the genomic landscape of two co-distributed woodpecker species whose population histories have been profoundly impacted by the Ice Age. We found that despite a dynamic demographic history, Downy and Hairy Woodpecker were able to maintain very large N_e_ even during glacial periods, which might have facilitated the action of natural selection. Supporting this conclusion, our results reveal a correlation between nucleotide diversity, recombination rate, and gene density, which suggests the effect of linked selection shaping the genomic landscape. In addition, we found that the magnitude of linked selection was associated with population-specific N_e_ trajectories, indicating that demography and natural selection operated in concert to shape patterns of polymorphism along the genome. This study adds to the growing body of literature supporting the role of natural selection in driving patterns of genome-wide variation but highlights the difficulty of interpreting the outcome of the interplay between genetic drift and natural selection in organisms with non-equilibrium demographic dynamics and large effective population sizes.

## Material and Methods

### Sample collection and whole genome sequencing

We collected 70 samples for both the Downy Woodpecker (*D. pubescens*) and Hairy Woodpecker (*D. villosus*) in each of seven populations (*n* = 10 per population) across their temperate North American ranges (Figure 1): New York (Northeast), Louisiana (Southeast), Minnesota (Midwest), New Mexico and Colorado (Southern Rockies), Wyoming (Northern Rockies), Washington (Pacific Northwest), and Alaska. The samples were obtained through museum loans of vouchered specimens and augmented by field collections in Wyoming, Louisiana, and Alaska (Table S1). We extracted genomic DNA from tissue samples using the MagAttract High Molecular Weight DNA Kit from Qiagen following manufacturer’s instructions (Qiagen, California, USA). These samples were then submitted for whole genome resequencing on a paired-end Illumina HiSeq X Ten machine at RAPiD Genomics (Gainesville, Florida, USA).

### Read alignment, variant calling and filtering

Raw reads were trimmed for Illumina adapters using Trimmomatic v0.36 [107] with the following parameters:” ILLUMINACLIP:TruSeq3-PE-2.fa:2:30:10:8:true”, resulting in an average of 35,689,979 paired reads per sample. Read quality was assessed with FastQC v0.11.4. [108]. Given the high synteny and evolutionary stasis of bird chromosomes [82], we produced a chromosome-length reference genome for Downy Woodpecker by ordering and orienting the scaffolds and contigs of the Downy Woodpecker genome assembly [52] along the 35 chromosomes of the Zebra finch (*Taeniopygia guttata*; version taeGut3.2.4) using Chromosemble from the Satsuma package [109]. We verified the completeness of this new reference by searching for a set of single-copy avian orthologs using BUSCO v2.0.1 (Benchmarking Universal Single-Copy Orthologs [110]). A total of 91.1% of these genes were present and complete in our pseudo-chromosome reference, indicating sufficient completeness. We finally transferred the prediction-based genome annotation of the Downy Woodpecker [52] by mapping the genomic coordinates of each annotated feature against the pseudo-chromosome reference using gmap [111]. A total of 99.98% of all the 14,443 annotated genes in Downy Woodpecker were successfully mapped to the pseudo-chromosome reference.

Trimmed reads for both Downy and Hairy Woodpecker were aligned against the pseudo-chromosome reference genome of the Downy Woodpecker using BWA v0.7.15 mem algorithm [112]. On average, 97.27% of reads from Downy Woodpecker and 96.38% of reads from Hairy woodpecker were successfully mapped, demonstrating that despite the large evolutionary distance between these two species [47], sequence conservation allows efficient mapping. Resulting sequence alignment/map (SAM) files were converted to their binary format (BAM) and sequence group information was added. Next, reads were sorted, marked for duplicates, and indexed using Picard (http://broadinstitute.github.io/picard/). The Genome Analysis Toolkit (GATK v3.6 [113]) was then used to perform local realignment of reads near insertion and deletion (indels) polymorphisms. We first used the RealignerTargetCreator tool to identify regions where realignment was needed, then produced a new set of realigned binary sequence alignment/map (BAM) files using IndelRealigner. The final quality of mapping was assessed using QualiMap v.2.2.1 [114].

We implemented two complementary approaches for the downstream analysis of genetic polymorphism. First, we used ANGSD v0.917 [53], a method that accounts for the genotype uncertainty inherent to low depth sequencing data by inferring genotype likelihoods instead of relying on genotype calls. We estimated genotype likelihoods from BAM files using the GATK model (*-GL 2* [113]*), retaining only sites present in at least 70% of sampled individuals (-minInd 50*) and with the following filters: a minimum mapping quality of 30 (*-minMapQ 30*), a minimum quality score of 20 (*-minQ* 20), a minimum frequency of the minor allele of 5% (*-minMaf 0*.*05*), and a P-value threshold for the allele-frequency likelihood ratio test statistic of 0.01 (*-SNP_pval 0*.*01*). Allele frequencies were estimated directly from genotype likelihoods assuming known major and minor alleles (*-doMajorMinor 1 -doMaf 1*). A total of 16,736,465 and 15,463,356 SNPs were identified for Downy and Hairy Woodpecker, respectively. Because several downstream analyses lack support for genotype likelihoods, we also called genotypes using GATK v3.8.0 [115]. First, we run HaplotypeCaller separately for each sample using the *--emitRefConfidence GVCF -minPruning 1 -minDanglingBranchLength 1* options to create one gVCF per individual, then we ran GenotypeGVCFs with default settings across all samples to jointly call genotypes. In the absence of a training SNP panel for our non-model species, we applied hard filtering recommendations from the Broad Institute’s Best Practices (https://gatk.broadinstitute.org/). We filtered SNPs with quality by depth below 2 (QD < 2.0), SNPs where reads with the alternative allele were shorter than those with the reference allele (ReadPosRankSum < -8), SNPs with evidence of strand bias (FS > 60.0 and SOR > 3.0), SNPs with root mean square of the mapping quality below 40 (MQ < 40.0), and SNPs in reads where the alternative allele had a lower mapping quality than the reference allele (MQRankSumTest < -12.5). In addition, we used VCFtools v0.1.17 [116] to retain only biallelic SNPs occurring in at least 75% of samples, with a minimal mean coverage of 2x, a maximum mean coverage of 100x, and a P-value above 0.01 for the exact test for Hardy-Weinberg Equilibrium. We applied three different minor allele frequency (maf) thresholds – 0.05 (for most analyses), 0.02 (for the estimation of recombination rates), and no threshold (for demographic analyses based on the SFS).

### Population structure

To assess population structure, we performed a principal components analysis (PCA) using the R package SNPRelate v3.3 [117]. We first applied the function *snpgdsLDpruning* to select a subset of unlinked SNPs (LD r^2^ threshold = 0.2), with < 25% missing data and a maf > 0.05, which resulted in a total of 71,228 SNPs for Downy Woodpecker and 71,763 SNPs for Hairy Woodpecker. We then used the function *snpgdsPCA* to calculate the eigenvectors and eigenvalues for the principal component analysis. We investigated population structure by looking at the first three principal components (PC1–PC3). In addition, we used NGSadmix [54], implemented in ANGSD [53], to investigate the number of genetic clusters, and associated admixture proportions for each individual. NGSadmix is a maximum likelihood approach analogous to STRUCTURE [118], but bases its inferences on genotype likelihoods instead of SNP calls, therefore accounting for the uncertainty of genotypes.

We also described the relationships among populations by building a maximum likelihood tree based on the polymorphism-aware phylogenetic model (PoMo [58]) implemented in IQ-Tree 2 [59]. PoMo is a phylogenetic method that accounts for incomplete lineage sorting inherent to population-level data by incorporating polymorphic states into DNA substitution models. We used a python script (https://github.com/pomo-dev/cflib) to convert our vcf files containing only intergenic SNPs into the input format of PoMo (counts file). IQ-Tree was run using the HKY+P model of sequence evolution with 100 non-parametric bootstraps to assess support. We used three samples from Hairy Woodpecker as an outgroup to root the tree for Downy Woodpecker, and vice versa.

We estimated pairwise F_ST_ values among populations in each species using ANGSD v0.917 [53]. We first produced site-allele-frequency likelihoods using the command *-doSaf*, followed by the *realSFS - fold 1* command to generate a folded site frequency spectrum (SFS). We then estimated weighted F_ST_ values using the *realSFS fst* command both globally and across non-overlapping 100 kb windows.

We investigated patterns of gene flow across the landscape using the estimated effective migration surface (EEMS [55]), which is a method to visualize variation in patterns of gene flow across a habitat. Low values of relative effective migration rate (*m*) indicate a rapid decay in genetic similarity in relation to geographic distances, which suggests the presence of barriers to gene flow. In contract, high values of *m* indicate larger genetic similarity than expected given the geographic distance, suggesting genetic connectivity. We generated pairwise identity-by-state (IBS) matrices using the -doIBS function in ANGSD [53] and used these matrices to represent dissimilarity between individuals. We ran EEMS using 200 demes and performed a single MCMC chain run with 1 ⨯ 10^7^ iterations following a burn-in of 5 ⨯ 10^6^, and a thinning of 9,999. We then checked the posterior probabilities to ensure convergence.

### Demographic inference

We inferred past changes in effective population size (N_e_) using Stairway Plot 2 [56], a method that leverages information contained in the site frequency spectrum (SFS) to estimate recent population history. Unlike methods based on the Sequentially Markov Coalescent (e.g, PSMC, SMC++), Stairway Plot 2 is applicable to a large sample of unphased whole genome sequences, and it is insensitive to read depth limitations. We estimated the folded site frequency spectrum for each population using the *realSFS* function in ANGSD [53]. For each population, we used the default 67% sites for training, and calculated median estimates and 95% pseudo-CI based on 200 replicates. We assumed a mutation rate of 4.007 × 10^−9^ mutations per site per generation, as estimated from coding regions of the Northern Flicker’s genome [57] and a generation time of one year for both species. We then utilized the estimates of N_e_ from Stairway Plot 2 across the past 500 kya to calculate the harmonic mean on linear-stepped time points, representing each population’s long-term N_e_.

We further investigated the demographic history of the two species using *fastsimcoal2* v2.6.0.3, a composite likelihood method that uses the joint site frequency spectrum (jSFS) to perform model selection and estimate demographic parameters [62]. We tested the support for two competing demographic models: (i) a model where all populations diverge synchronously from a single large refugium and expand independently with asymmetric gene flow, and (ii) a bifurcating model where populations diverge at different times from multiple refugia and expand independently with asymmetric gene flow. Since we only need a reasonably large subset of the genome to get an accurate estimate of the site frequency spectrum [119], we generated the four-population folded jSFS from a set of high quality SNPs with no maf filtering (Downy: 6,030,759 SNPs; Hairy: 7,967,215 SNPs) present in chromosome 1 using easySFS.py (https://github.com/isaacovercast/easySFS). We projected the jSFS down to 20 chromosomes (i.e., 10 diploid samples) per population to avoid issues associated with differences in sample size and missing data. To minimize the impact of selection, we only included sites in non-coding regions of the genome. All models followed the topology of the population tree obtained from IQ-Tree 2 and assumed a mutation rate of 4.007 ⨯ 10^−9^ mutations per site per generation. For each model, we conducted 75 iterations of the optimization procedure, each with 40 expectation conditional maximization cycles and 100,000 genealogical simulations per cycle. We performed model selection using the run with the highest likelihood for each model. For each species, we chose the model with the largest relative Akaike information criterion (AIC_w_) as the best-fit model. We obtained 95% pseudo-CI for parameter estimates by performing 100 parametric bootstrap estimates simulating jSFSs under the best model and re-estimating parameters using these simulated datasets.

### Genetic diversity, recombination rates, and linkage disequilibrium

We compared genetic diversity among populations of the two species by estimating the genome-wide pairwise nucleotide diversity θ_π_ and the Watterson estimator of the rescaled mutation rate per base θ_W_ sing ANGSD [53]. We first ran the command *-doSaf* 1 *-minMapQ 30 -minQ 20* in ANGSD to generate site-allele-frequency likelihoods based on the GATK model [115], then we used -realSFS with the option *-fold 1* to estimate the folded SFS. ANGSD was also used to estimate genome-wide Tajima’s D. We estimated recombination rates (*r* = recombination rate per base pair per generation) along the genome of the two species using ReLERNN, a deep learning algorithm [63]. ReLERNN takes as input a vcf file and simulates training, validation, and test datasets matching the empirical distribution of θ_W_. ReLERNN then uses the raw genotype matrix and a vector of genomic coordinates to train a model that directly predicts per-base recombination rates (as opposed to a population-scaled recombination rate) across sliding windows [63]. To reduce the impact of population structure on estimates, we restricted the prediction of recombination rates to the Eastern populations (Northeast + Southeast + Midwest), the genetic cluster with most samples. Given the conserved landscape of recombination in birds, we do not expect major differences in recombination across populations [82]. We used the SNP dataset with maf > 0.02 and ran the analysis with default settings. Because ReLERNN is robust to demographic model misspecification [63], we simulated an equilibrium model considering a mutation rate of 4.007 × 10^−9^ mutations per generation [57] and assuming a generation time of one year. Finally, we explored the recombination history of each population by analyzing their patterns of linkage disequilibrium (LD) decay using PopLDdecay [120]. We calculated pairwise D’/r^2^ using the default maximum distance between SNPs of 300 kb and plotted it as a function of genomic distance (in kb).

### Genomic predictors of regional variation in nucleotide diversity

To investigate the factors shaping the genomic landscape of diversity in the two woodpecker species, we tested the effect of (i) recombination rate, (ii) gene density, and (iii) GC content on regional patterns of nucleotide diversity. We computed pairwise nucleotide diversity (θ_π_) across 100 kb non-overlapping windows using ANGSD [53]. We first used the *-doThetas* function to estimate the site-specific nucleotide diversity from the posterior probability of allele frequency (SAF) using the estimated site frequency spectrum (SFS) as a prior. Then, we ran the *thetaStat do_stat* command to perform the sliding windows analysis. To quantify variation in recombination rates, we calculated weighted averages of recombination rates estimated in ReLERNN across 100 kb non-overlapping windows. We assessed gene density (i.e., density of targets of selection) as the proportion of coding sequence (in number of base-pairs) for any given 100 kb non-overlapping window and estimated GC content in each 100 kb non-overlapping window using the function *GC* of the R package *seqinr* version 3.6-1 [121]. We fit a general linear regression in R to assess the relationship between nucleotide diversity (θ_π_) and the three predictor variables – recombination rate, gene density, and base composition. We also fit a LOESS model to account for the potential non-linearity of these relationships using the R package *caret* [122]. Models were trained using cross-validation of 80% of the total data. To control for the collinearity among these variables, we also ran a principal component regression (PCR). PCR is a technique that summarizes the predictor variables into orthogonal components (PCs) before performing regression, therefore removing the correlation among variables. PCR was conducted using the R package pls [123]. All variables were Z-transformed before these analyses.

We also investigated the association between patterns of intraspecific population differentiation (F_ST_) and intrinsic properties of the genome (i.e., nucleotide diversity and recombination rates). To summarize the genomic landscape of differentiation into a single response variable we employed two approaches: for each 100 kb windows, we (i) calculated the average F_ST_ across all pairwise population comparisons; (ii) we performed a principal component analysis and extracted that first principal component (PC1) that explained the greatest covariance among all pairwise population comparisons.

### Natural selection and genetic load

To estimate the genetic load of each species and populations, we first used the software snpEff v4.1 [124] to classify SNPs into one of four categories of functional impact, according to the predicted effect of the gene annotation – (i) modifiers: variants in non-coding regions of the genome (e.g, introns, intergenic) whose effects are hard to predict; (ii) low: variants in coding sequences that cause no change in amino acid (i.e., synonymous); (iii) moderate: variants in coding sequences that cause a change in amino acid (i.e., nonsynonymous); and (iv) high: variants in coding sequences that cause gain or loss of start and stop codon. We then selected a subset of 12 individuals with the lowest percentage of missing data (therefore, maximizing the total number of sites) in each species to polarize our SNPs. To do so, we looked for biallelic SNPs in Downy Woodpecker for which one of the alleles were fixed in Hairy Woodpecker and vice versa. The allele fixed in the outgroup was assumed to be the ancestral state. This is a sensitive step in the estimation of genetic load, so we only kept SNPs for which the ancestral state could be determined unambiguously [70]. We ended up with a total set of 363,903 polarized SNPs across the genome.

We characterized the site frequency spectrum (SFS) for each type of variant (according to the impact inferred from snpEff) by estimating the total frequency of each derived allele and calculating the proportion of each allele frequency bin. As a proxy for genetic load, for each individual, we estimated the ratio of the number of derived alleles of high impact (i.e., loss of function) in homozygosity over the number of derived alleles of low impact (i.e, synonymous) in homozygosity. This metric assumes a recessive model, in which derived alleles are only deleterious when in a homozygous state. We therefore also considered an additive model (i.e, semi-dominant) that assumes that derived alleles have deleterious effects in both homozygosity and heterozygosity. For this metric, we counted the total number of derived alleles, instead of only the ones in homozygosity [70].

To look at selection over a deeper evolutionary scale, we estimated dN/dS, the ratio of nonsynonymous over synonymous substitution, using a set of 397 genes that were orthologous across Downy Woodpecker, Hairy Woodpecker and two avian outgroups – Chicken (*Gallus gallus*) and Zebra Finch (*Taeniopygia guttata*). We identified orthologous genes across all four species using the software JustOrthlogs [125] and only kept well-aligned loci. We first downloaded Ensembl genome assemblies and gene annotations for version *GRCg6a* and *bTaeGut1_v1*.*p* of the Chicken and Zebra Finch genome, respectively (Ensembl v103). We then extracted coding sequences (CDS) for all identified orthologs from their respective reference genomes using a GFF3 parser included in JustOrthologs and aligned them with the frameshift-aware MACSE software [126]. We used the parameter setting *--min_percent_NT_at_ends 0*.*3* and *-codonForInternalStop NNN* for aligning and exporting sequences. The resulting amino-acid alignments were inspected with HMMcleaner to mask sites that were likely misaligned. We finally used *codeml* to estimate the overall dN/dS ratio along each branch of the tree assuming a one-ratio branch model in PAML [127].

## Acknowledgments

We thank the following individuals and institutions for providing tissue samples and specimen loans for this study: A. G. Navarro Sigüenza (Museo de Zoología; UNAM), J. Klicka/S. Birk/R. Faucett (University of Washington Burke Museum), C. M. Milensky (Smithsonian Institution), C. Dardia (Cornell University Museum of Vertebrates), G. Spellman/A. Doll (Denver Museum of Nature & Science), B. Marks/S. Hackett/J. Bates (Field Museum of Natural History), F. Sheldon/D. Dittmann (LSU Museum of Natural Science), K. Barker/T. Imfeld (Midwest Museum of Natural History), C. Witt/A. Johnson/M. Anderson (Museum of Southwestern Biology). This work would not be possible without the assistance of Lucas DeCicco, Matt Brady, and Paul Sweet, who were very generous to help collect samples in the field. We thank Kaiya Provost, Jon Merwin, Vivien Chua, Glenn Seehozer, Gregory Thom, Elkin Tenorio, William Mauck, Lukas Musher, Laís Coelho, Amanda Rocha, Bruno Almeida, Isaac Overcast, Joel Cracraft, Frank Burbrink, Molly Przeworski, Deren Eaton, and Don Melnick for their invaluable input during the development and writing of this manuscript.

## Author contributions

This study was conceived and designed by L.R.M. and B.T.S. A subset of samples was collected and made available by J.K. L.R.M. conducted all wet lab, bioinformatic analyses, and drafted the paper with input from all authors.

## Data and materials availability

Whole-genome resequencing data are available at the Sequence Read Archive (https://ncbi.nlm.nih.gov/sra) under accession numbers #. All data needed to evaluate the conclusions in the paper are present in the paper and/or the Supplementary Materials. Code used in this study is avaxsilable at https://github.com/lucasrocmoreira/Moreira-et-al-2022. Additional data related to this paper may be requested from the authors.

