## Supplementary Materials for "Demography and linked selection interact to shape the genomic landscape of codistributed woodpeckers during the Ice Age"

**This PDF file includes:**

Supplementary Text
Figs. S1 to S6
Tables S1 to S3

**Supplementary Text**

*Demographic dynamics*

We inferred a dynamic demographic history for Downy and Hairy Woodpecker from Stairway Plot 2 [1]. At around 500 kya, both Downy and Hairy Woodpecker had an Ne of nearly 300,000 individuals and population sizes dropped between 500 kya and 300 kya to approximately 60,000 individuals. This was followed by an episode of demographic expansion when populations increased nearly 10-fold. A second population decline occurred around the onset of the Last Glacial Period (LGP; 115 kya), but the exact timing varied across populations. A final spike in Ne occurred after the Last Glacial Maximum (LGM; 21 kya), when a more than 20-fold population growth occurred in the East, likely a result of the glacial retreat. In the Rocky Mountains, a final explosive expansion preceded the LGM, whereas in Alaska, a final population expansion occurred more recently in the past 3–15 kya.

*Genetic diversity*

The broadly distributed Downy and Hairy Woodpeckers exhibited genome-wide levels of nucleotide diversity larger than those observed in most bird species [2–6]. Mean values of nucleotide diversity were slightly larger in Hairy ($\theta\pi$= 0.0064; within population = 0.0045–0.0065) than in Downy Woodpecker ($\theta\pi$= 0.006; within population = 0.0049–0.0061). Genetic diversity was lowest in Alaska (AK; $\theta\pi$ Downy= 0.0049, $\theta\pi$ Hairy = 0.0045) and largest in the Northern Rockies (NR; $\theta\pi$ Downy= 0.0061, $\theta\pi$ Hairy = 0.0065) and Southeast (SE; $\theta\pi$ Downy= 0.0059, $\theta\pi$ Hairy = 0.0062; Figure 4a). Regionally, levels of genetic diversity in populations of Downy Woodpecker surpassed all of those of Hairy Woodpecker, with exception of Northern Rockies and Southeast (Figure 3b). Genome-wide values of Tajima's D [7] were consistently negative across six populations of Downy and Hairy Woodpecker (Downy: -1.35– -0.39; Hairy: -0.88– -24; Figure 3b). Negative values of genome-wide Tajima's D indicate an excess of low frequency alleles and are suggestive of population expansion. In Alaska, however, genome-wide Tajima's D were positive (Downy: 0.19; Hairy: 0.42), indicating ongoing population contraction or very recent population expansion [8].

*Population structure and demographic history*

We found that population structure was spatially congruent between Downy and Hairy Woodpecker, which is likely driven by drift and varying gene flow regimes across the landscape. Both species are characterized by four genetic clusters that are consistent with previous phylogeographic studies – East, Alaska, Rocky Mountains, and Pacific Northwest [9–11]. Genetic structure in Hairy Woodpecker shows a clear east-west subdivision, which is estimated to have occurred in the Mid-Pleistocene (513–561 kya) when glacial-interglacial cycles increased in length and intensity [12]. An east-west split is a common biogeographic pattern observed in widely distributed North American birds [13–16]. Our demographic analyses supported the existence of at least two glacial refugia that isolated populations of Hairy Woodpecker on either side of North America and gave origin to the four genetic clusters. Previous paleoclimate modelling supports multiple southern refugia during the LGM [9–11].

Despite geographic congruence, genetic structure in Downy Woodpecker shows a few differences. First, we found genome-wide population differentiation to be higher in Hairy Woodpecker (average FST = 0.1; 0.03–0.19) than in Downy Woodpecker (average FST = 0.08; 0.03–0.16). These results agree with previous genetic studies using a smaller number of loci, which reported very shallow population differentiation in Downy Woodpecker [11,17].

However, it appears that genetic diversity and structure in the mtDNA of Downy Woodpecker is much lower than in the nuclear genome [11,17]. Such discrepancies may reflect inherent differences in Ne between different genomes, a possible selective sweep that could have reduced mtDNA diversity, or even sex-biased dispersal [18]. In fact, females of Downy Woodpecker have a higher tendency for long-distance dispersal than males, which in part could explain the homogeneity of the mitochondrial genome [19]. Regardless, an elevated FST in Hairy Woodpecker when compared to Downy Woodpecker indicates that gene flow in Hairy Woodpecker might be more restricted. Second, the clear east-west subdivision observed in Hairy Woodpecker was not seen in Downy Woodpecker. Our phylogenetic tree shows that Alaska was the first population to diverge from the clade containing all other populations, followed by the Rocky Mountains. Such a topology may arise if Alaska contained suitable habitat for populations to persist through glacial cycles, which seems to have been the case for several boreal species [20–22]. Nevertheless, the low genetic diversity and signature of recent population expansion in Alaska favors a scenario of colonization [11]. Moreover, we found support for a model in which all daughter populations of Downy Woodpecker arise simultaneously from a single ancestral population (i.e, polytomy). Under this scenario, the elevated differentiation of Alaska could be due to its further distance from other populations.

Our demographic analyses reveal a dynamic population history for Hairy and Downy Woodpecker during the Ice Age. Both species underwent repeated cycles of population contraction and expansion, consistent with the climatic fluctuations of the Pleistocene. Two main episodes of bottleneck followed by expansion can be detected in our dataset – the first one occurring in the Mid Pleistocene, between 500 and 300 kya, when range contraction and persistent isolation in glacial refugia have likely contributed to population differentiation. The second one occurred during and after the Last Glacial Period (LGP; 70 kya – 8 kya), when populations underwent strong decline followed by a more than 20-fold growth. The timing and magnitude of these changes differed across geographic regions. Despite strong variation in Ne over the past 500 ky, our data indicates that Downy and Hairy Woodpecker have been resilient enough to maintain relatively large populations, which favored the maintenance of very high genetic diversity, even in the face of repeated bottlenecks. It is worth noting, however, that estimates of Ne are critically dependent on the choice of mutation rate and generation time, which are uncertain parameters.

*GC content and genetic diversity*

We observed an association between nucleotide diversity ($\theta\pi$) and GC content in Downy and Hairy Woodpecker. The frequency of GC nucleotides is known to be considerably higher in coding sequence when compared to noncoding [23]. Accordingly, we found a significant correlation between GC content and gene density, but our principal component regression failed to dissect the effect of these two variables on patterns of nucleotide diversity. We also found a weak correlation between GC content and recombination rate in Hairy Woodpecker. A mechanism widely postulated to explain this correlation is GC-biased gene conversion (gBGC), a process whereby AT/GC heterozygotes are more likely to pass GC nucleotides to descendants during meiotic recombination [24]. This mechanism mimics selection favoring GC and is tightly linked to variation in recombination rate, so that high GC content is expected in regions of high recombination. In birds, gBGC is an important factor shaping GC content along the genome and is positively correlated with divergence at neutral sites [25].

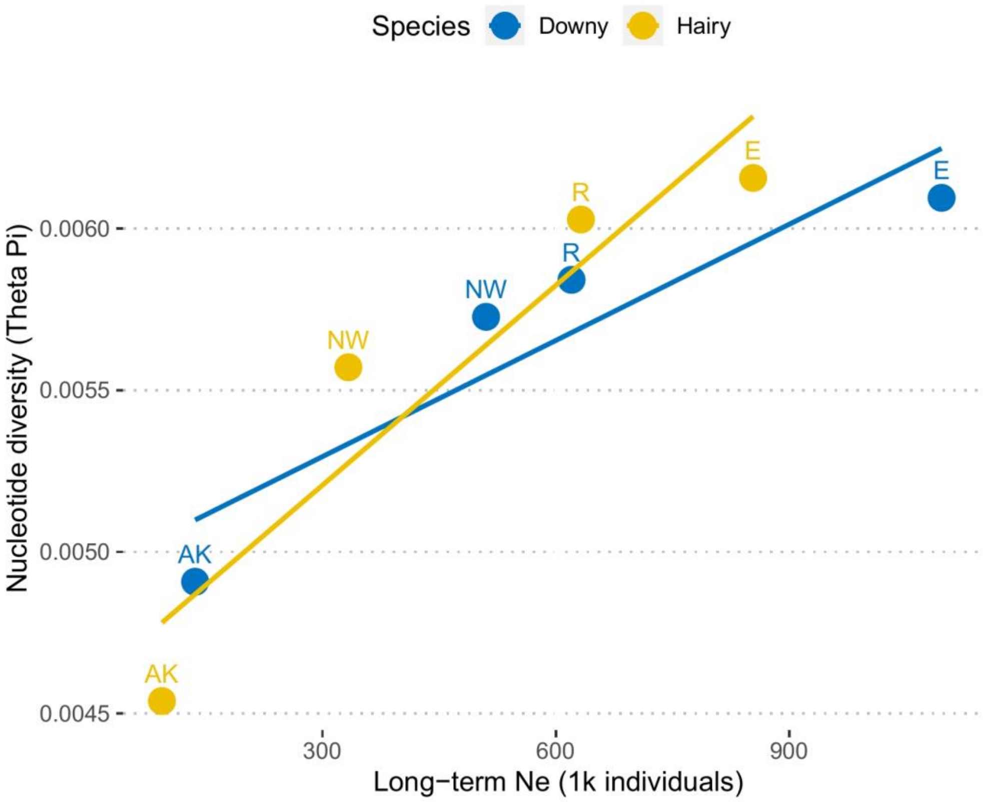


**Figure S1. Relationship between pairwise nucleotide diversity ($\theta_\pi$) and long-term effective population size ($N_e$).**

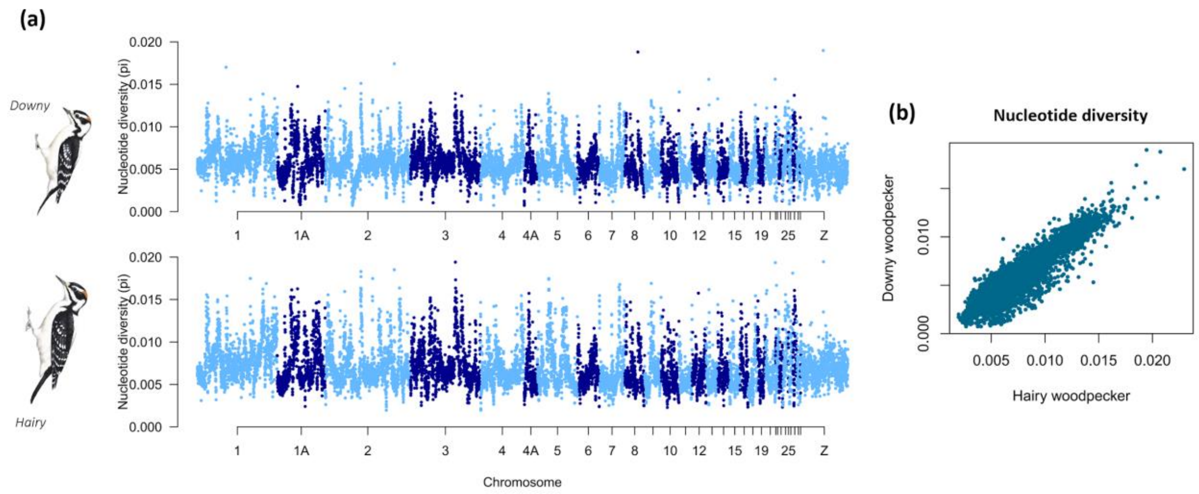


**Figure S2. Correlated landscape of diversity in Downy (top) and Hairy (bottom) Woodpecker. (a)** Manhattan plot of nucleotide diversity ($\theta_\pi$) along the genome. Each point represents a non-overlapping 100 kb window. Colors depict different chromosomes. **(b)** Scatterplot of the correlation in nucleotide diversity between Downy and Hairy Woodpecker. Illustrations reproduced with permission from Lynx Edicions.

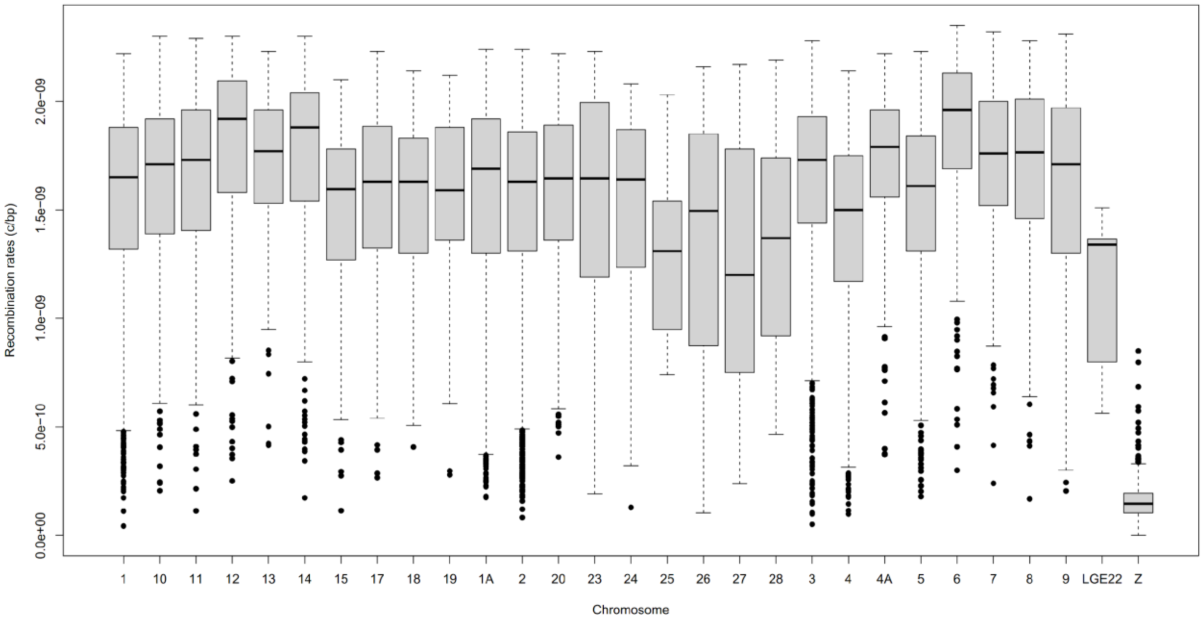


**Figure S3. Boxplot of recombination rate in each chromosome of Downy Woodpecker.** Horizontal lines indicate medians, boxes span the interquartile range (IQR), and points represent outliers.

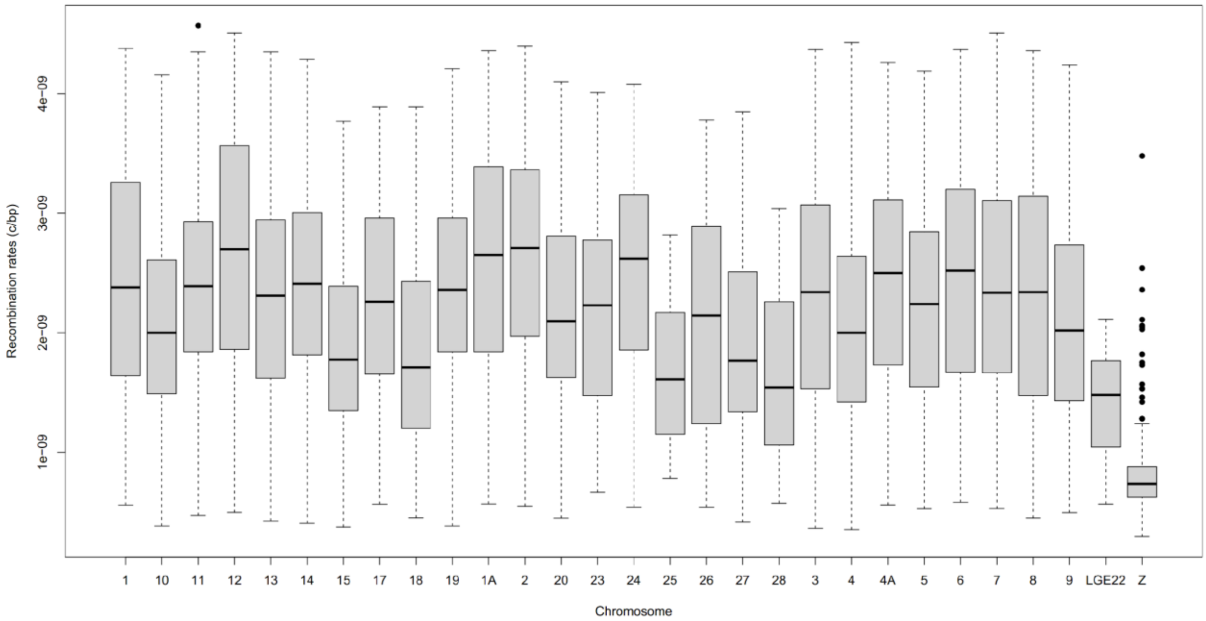


**Figure S4. Boxplot of recombination rate in each chromosome of Hairy Woodpecker.** Horizontal lines indicate medians, boxes span the interquartile range (IQR), and points represent outliers.

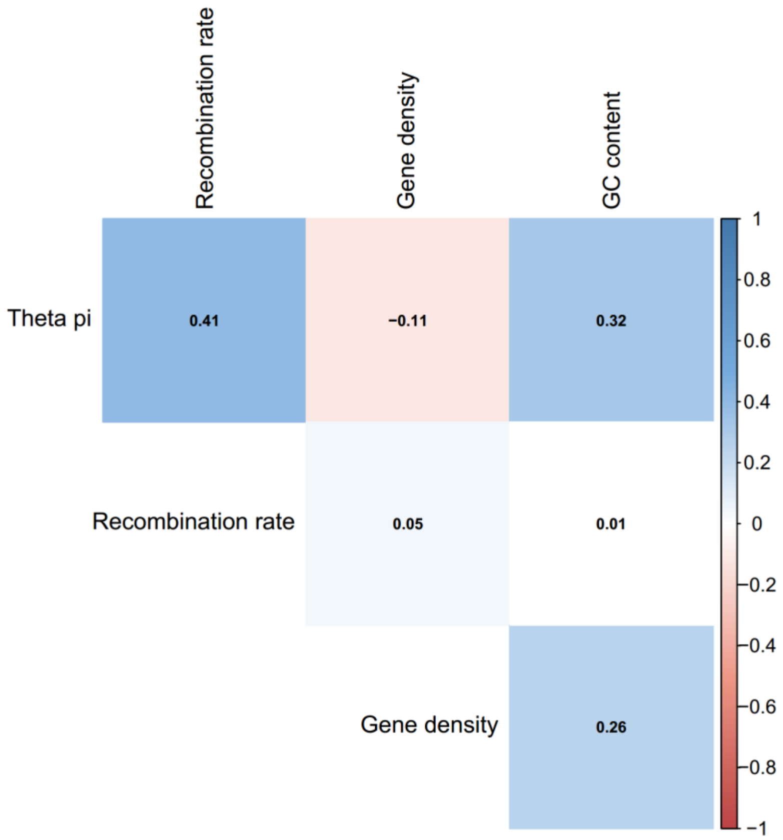


**Figure S5. Correlation among genomic variables in Downy Woodpecker.** Colder colors represent positive values of Pearson’s r. Warmer colors represent negative values of Pearson’s r.

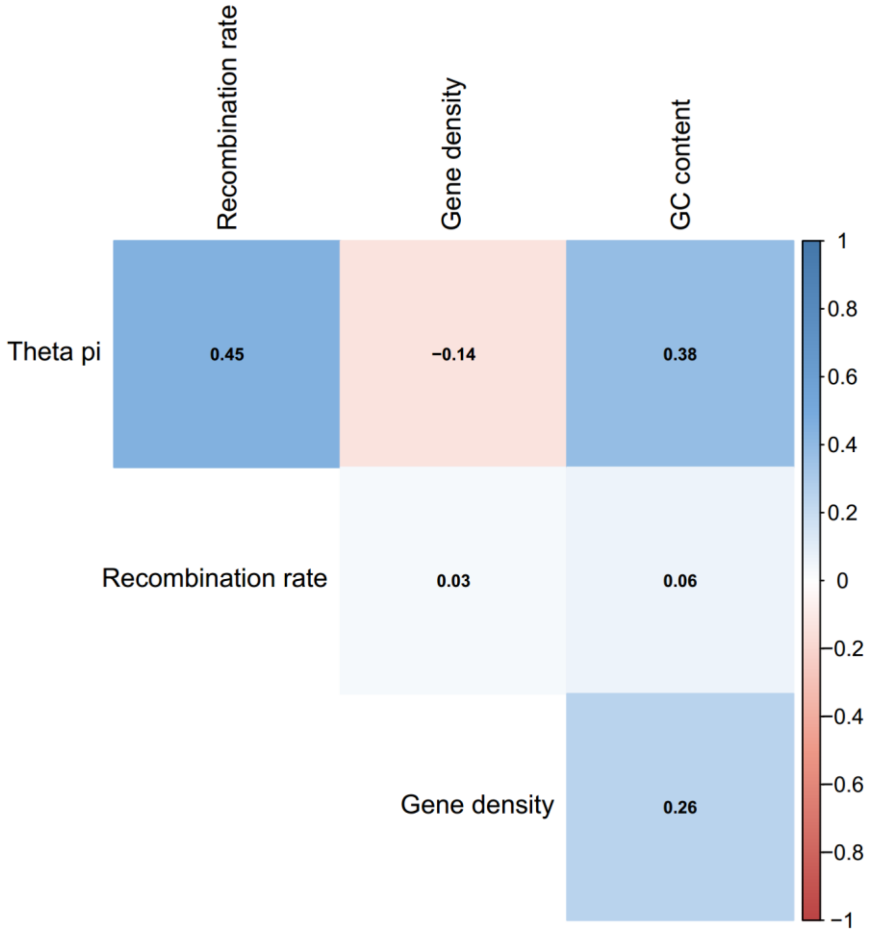


**Figure S6. Correlation among genomic variables in Hairy Woodpecker.** Colder colors represent positive values of Pearson's r. Warmer colors represent negative values of Pearson's r.

**Table S1. Sample information.**

| Species | Sample ID | Population | Institution | Voucher Number | Sex |
|---|---|---|---|---|---|
| *D. pubescens* | PP-NE-26 | NE | AMNH | DOT 17125 | F |
| *D. pubescens* | PP-NE-37 | NE | AMNH | DOT 20863 | F |
| *D. pubescens* | PP-NE-38 | NE | AMNH | DOT 20864 | F |
| *D. pubescens* | PP-NE-39 | NE | AMNH | DOT 20865 | F |
| *D. pubescens* | PP-NE-40 | NE | AMNH | DOT 20866 | F |
| *D. pubescens* | PP-NE-42 | NE | AMNH | DOT 21195 | M |
| *D. pubescens* | PP-NE-43 | NE | AMNH | DOT 21196 | F |
| *D. pubescens* | PP-NE-47 | NE | CUMV | 55468 | F |
| *D. pubescens* | PP-NE-48 | NE | CUMV | 55567 | F |
| *D. pubescens* | PP-NE-49 | NE | CUMV | 55647 | M |
| *D. pubescens* | PP-MW-2 | MW | FMNH | 442445 | M |
| *D. pubescens* | PP-MW-3 | MW | FMNH | 442446 | M |
| *D. pubescens* | PP-MW-4 | MW | FMNH | 442447 | M |
| *D. pubescens* | PP-MW-7 | MW | FMNH | 461726 | M |
| *D. pubescens* | PP-MW-11 | MW | FMNH | 485368 | F |
| *D. pubescens* | PP-MW-18 | MW | MMNH | 47208 | F |
| *D. pubescens* | PP-MW-19 | MW | MMNH | 47209 | M |
| *D. pubescens* | PP-MW-20 | MW | MMNH | 47292 | M |
| *D. pubescens* | PP-MW-21 | MW | MMNH | 47293 | F |
| *D. pubescens* | PP-MW-22 | MW | MMNH | 49295 | M |
| *D. pubescens* | PP-SR-9 | SR | DMNS | 44330 | M |
| *D. pubescens* | PP-SR-10 | SR | DMNS | 44404 | F |
| *D. pubescens* | PP-SR-12 | SR | MSB | 26384 | M |
| *D. pubescens* | PP-SR-13 | SR | MSB | 26652 | M |
| *D. pubescens* | PP-SR-15 | SR | MSB | 28961 | F |
| *D. pubescens* | PP-SR-16 | SR | MSB | 29394 | F |
| *D. pubescens* | PP-SR-17 | SR | MSB | 29766 | F |
| *D. pubescens* | PP-SR-18 | SR | MSB | 30563 | F |
| *D. pubescens* | PP-SR-19 | SR | MSB | 30599 | M |
| *D. pubescens* | PP-SR-21 | SR | MSB | 41048 | M |
| *D. pubescens* | PP-NW-5 | NW | UWBM | 79382 | F |
| *D. pubescens* | PP-NW-8 | NW | UWBM | 81721 | F |
| *D. pubescens* | PP-NW-9 | NW | UWBM | 85226 | M |
| *D. pubescens* | PP-NW-10 | NW | UWBM | 85940 | F |
| *D. pubescens* | PP-NW-11 | NW | UWBM | 89835 | F |
| *D. pubescens* | PP-NW-12 | NW | UWBM | 91432 | F |
| *D. pubescens* | PP-NW-13 | NW | UWBM | 119080 | M |
| *D. pubescens* | PP-NW-15 | NW | UWBM | 121308 | M |
| *D. pubescens* | PP-NW-16 | NW | UWBM | 121544 | F |
| *D. pubescens* | PP-NW-18 | NW | UWBM | 122655 | F |
| *D. pubescens* | PP-AK-1 | AK | AMNH | LRM10 | M |
| *D. pubescens* | PP-AK-2 | AK | AMNH | LRM46 | F |
| *D. pubescens* | PP-AK-3 | AK | AMNH | LRM50 | F |
| *D. pubescens* | PP-AK-4 | AK | AMNH | LRM49 | F |
| *D. pubescens* | PP-AK-5 | AK | AMNH | LRM52 | F |

| | | | | | |
|---|---|---|---|---|---|
| *D. pubescens* | PP-AK-6 | AK | AMNH | LRM51 | F |
| *D. pubescens* | PP-AK-7 | AK | AMNH | LRM47 | M |
| *D. pubescens* | PP-AK-8 | AK | AMNH | LRM48 | F |
| *D. pubescens* | PP-AK-9 | AK | AMNH | LRM53 | M |
| *D. pubescens* | PP-AK-10 | AK | AMNH | LRM74 | F |
| *D. pubescens* | PP-NR-01 | NR | AMNH | LRM054 | F |
| *D. pubescens* | PP-NR-02 | NR | AMNH | LRM058 | F |
| *D. pubescens* | PP-NR-03 | NR | AMNH | LRM059 | M |
| *D. pubescens* | PP-NR-04 | NR | AMNH | LRM067 | M |
| *D. pubescens* | PP-NR-05 | NR | AMNH | LRM068 | M |
| *D. pubescens* | PP-NR-06 | NR | AMNH | LRM069 | F |
| *D. pubescens* | PP-NR-07 | NR | AMNH | LRM070 | M |
| *D. pubescens* | PP-NR-08 | NR | AMNH | LRM071 | M |
| *D. pubescens* | PP-NR-09 | NR | AMNH | LRM072 | F |
| *D. pubescens* | PP-NR-10 | NR | AMNH | LRM073 | F |
| *D. pubescens* | PP-SE-01 | SE | LSUMZ | B45645 | F |
| *D. pubescens* | PP-SE-02 | SE | LSUMZ | B45652 | |
| *D. pubescens* | PP-SE-08 | SE | LSUMZ | B56712 | M |
| *D. pubescens* | PP-SE-09 | SE | LSUMZ | B56924 | F |
| *D. pubescens* | PP-SE-10 | SE | LSUMZ | B56926 | M |
| *D. pubescens* | PP-SE-12 | SE | LSUMZ | B57578 | |
| *D. pubescens* | PP-SE-14 | SE | LSUMZ | B62479 | |
| *D. pubescens* | PP-SE-15 | SE | LSUMZ | B62481 | |
| *D. pubescens* | PP-SE-16 | SE | UWBM | 96589 | F |
| *D. pubescens* | PP-SE-18 | SE | UWBM | 105426 | F |
| *D. villosus* | PV-NE-01 | NE | AMNH | | F |
| *D. villosus* | PV-NE-27 | NE | AMNH | DOT 18646 | M |
| *D. villosus* | PV-NE-28 | NE | AMNH | DOT 18681 | M |
| *D. villosus* | PV-NE-29 | NE | AMNH | DOT 18682 | F |
| *D. villosus* | PV-NE-31 | NE | AMNH | DOT 18789 | M |
| *D. villosus* | PV-NE-32 | NE | AMNH | DOT 20174 | M |
| *D. villosus* | PV-NE-36 | NE | AMNH | DOT 21148 | M |
| *D. villosus* | PV-NE-37 | NE | AMNH | DOT 21149 | F |
| *D. villosus* | PV-NE-47 | NE | AMNH | DOT 22674 | M |
| *D. villosus* | PV-NE-51 | NE | AMNH | DOT 23032 | F |
| *D. villosus* | PV-NW-8 | NW | MMNH | 47181 | F |
| *D. villosus* | PV-NW-12 | NW | MMNH | 47215 | F |
| *D. villosus* | PV-NW-16 | NW | UWBM | 49955 | F |
| *D. villosus* | PV-NW-17 | NW | UWBM | 49958 | F |
| *D. villosus* | PV-NW-18 | NW | UWBM | 62606 | F |
| *D. villosus* | PV-NW-21 | NW | UWBM | 79778 | M |
| *D. villosus* | PV-NW-23 | NW | UWBM | 84259 | F |
| *D. villosus* | PV-NW-25 | NW | UWBM | 89951 | M |
| *D. villosus* | PV-NW-26 | NW | UWBM | 109529 | M |
| *D. villosus* | PV-NW-27 | NW | UWBM | 109530 | F |
| *D. villosus* | PV-SE-1 | SE | LSUMZ | 804 | M |
| *D. villosus* | PV-SE-2 | SE | LSUMZ | 3840 | M |
| *D. villosus* | PV-SE-3 | SE | LSUMZ | 8532 | F |

| | | | | | |
|---|---|---|---|---|---|
| *D. villosus* | PV-SE-4 | SE | UWBM | 116288 | F |
| *D. villosus* | PV-SE-5 | SE | UWBM | 116289 | F |
| *D. villosus* | PV-SE-6 | SE | AMNH | LRM86 | M |
| *D. villosus* | PV-SE-7 | SE | AMNH | LRM87 | M |
| *D. villosus* | PV-SE-8 | SE | AMNH | LRM88 | M |
| *D. villosus* | PV-SE-9 | SE | AMNH | LRM89 | F |
| *D. villosus* | PV-SE-10 | SE | AMNH | LRM90 | F |
| *D. villosus* | PV-MW-2 | MW | FMNH | 387974 | M |
| *D. villosus* | PV-MW-3 | MW | FMNH | 432747 | F |
| *D. villosus* | PV-MW-7 | MW | FMNH | 477418 | F |
| *D. villosus* | PV-MW-9 | MW | FMNH | 480382 | M |
| *D. villosus* | PV-MW-11 | MW | FMNH | 486061 | F |
| *D. villosus* | PV-MW-12 | MW | FMNH | 487467 | F |
| *D. villosus* | PV-MW-13 | MW | MMNH | 43391 | F |
| *D. villosus* | PV-MW-14 | MW | MMNH | 43570 | M |
| *D. villosus* | PV-MW-15 | MW | MMNH | 47218 | M |
| *D. villosus* | PV-MW-16 | MW | MMNH | 47646 | F |
| *D. villosus* | PV-SR-6 | SR | DMNS | 43511 | |
| *D. villosus* | PV-SR-12 | SR | DMNS | 46730 | M |
| *D. villosus* | PV-SR-16 | SR | DMNS | 47414 | M |
| *D. villosus* | PV-SR-18 | SR | MSB | 26715 | F |
| *D. villosus* | PV-SR-19 | SR | MSB | 29251 | F |
| *D. villosus* | PV-SR-23 | SR | MSB | 39737 | F |
| *D. villosus* | PV-SR-24 | SR | MSB | 40436 | M |
| *D. villosus* | PV-SR-27 | SR | MSB | 40790 | F |
| *D. villosus* | PV-SR-30 | SR | MSB | 45060 | F |
| *D. villosus* | PV-SR-31 | SR | MSB | 45150 | F |
| *D. villosus* | PV-AK-1 | AK | AMNH | LHD1133 | M |
| *D. villosus* | PV-AK-2 | AK | AMNH | LHD1134 | F |
| *D. villosus* | PV-AK-3 | AK | AMNH | LHD1137 | M |
| *D. villosus* | PV-AK-4 | AK | AMNH | LHD1106 | F |
| *D. villosus* | PV-AK-5 | AK | AMNH | LHD1105 | M |
| *D. villosus* | PV-AK-6 | AK | AMNH | LHD1107 | M |
| *D. villosus* | PV-AK-7 | AK | AMNH | LHD1108 | F |
| *D. villosus* | PV-AK-8 | AK | AMNH | LHD1117 | F |
| *D. villosus* | PV-AK-9 | AK | AMNH | LHD1118 | M |
| *D. villosus* | PV-AK-10 | AK | AMNH | LHD1138 | F |
| *D. villosus* | PV-NR-1 | NR | AMNH | LRM055 | F |
| *D. villosus* | PV-NR-2 | NR | AMNH | LRM056 | F |
| *D. villosus* | PV-NR-3 | NR | AMNH | LRM057 | M |
| *D. villosus* | PV-NR-4 | NR | AMNH | LRM060 | M |
| *D. villosus* | PV-NR-5 | NR | AMNH | LRM061 | F |
| *D. villosus* | PV-NR-6 | NR | AMNH | LRM062 | F |
| *D. villosus* | PV-NR-7 | NR | AMNH | LRM063 | M |
| *D. villosus* | PV-NR-8 | NR | AMNH | LRM064 | |
| *D. villosus* | PV-NR-9 | NR | AMNH | LRM065 | F |
| D. villosus | PV-NR-10 | NR | AMNH | LRM066 | M |

NE: Northeast; SE: Southeast; MW: Mid-West; SR: Southern Rockies; NR: Northern Rockies; NW: Pacific Northwest; AK: Alaska. F: female, M: male. AMNH: American Museum of Natural History; CUMV: Cornell University Museum of Vertebrates. FMNH: Field Museum of Natural History; MMNH: Midwest Museum of Natural History; DMNS: Denver Museum of Nature & Sciences, MSB: Museum of Southwestern Biology; UWBM: University of Washington Burke Museum; LSUMZ: Louisiana State University Museum of Zoology.

**Table S2. Model selection in *fastsimcoal2*.**

| Species | Model | Max ln(L) | Number of parameters | AIC | Relative weight |
|---|---|---|---|---|---|
| Downy Woodpecker | Single ancestral population | -4361743.523 | 26 | 8723539 | 1 |
| | Two ancestral populations | -4365454.312 | 30 | 8730969 | 0 |
| Hairy Woodpecker | Single ancestral population | -3572642.346 | 26 | 7145337 | 0 |
| | Two ancestral populations | -3571022.1 | 30 | 7142104 | 1 |

**Table S3. Parameter estimates for the best model in *fastsimcoal2* and their respective 95% confidence intervals.**

| Species | | Ne_AK | Ne_E | Ne_R | Ne_NW | B-Ne_AK | B-Ne_E | B-Ne_R | B-Ne-NW |
|---|---|---|---|---|---|---|---|---|---|
| Downy Woodpecker | Estimate | 1908367 | 8185752 | 9765060 | 1744261 | 533967 | 454210 | 204653 | 1556039 |
| | Lower 95% CI | 1016566 | 1046137 | 1015006 | 1015893 | 4.55E+02 | 8.32E+02 | 0 | 7.99E+02 |
| | Upper 95% CI | 3003138 | 11273989 | 35811351 | 2675784 | 2.23E+07 | 4426606 | 1782097 | 1.45E+08 |
| | | **Anc_Ne** | **T_div** | **T_div_E** | **T_div_W** | **T_exp_AK** | **T_exp_E** | **T_exp_R** | **T_exp_NW** |
| | Estimate | 7964011 | 516553 | 251877 | 252967 | 383864 | 309059 | 2.53E-06 | 2.15E-07 |
| | Lower 95% CI | 5653252 | 241530 | 114423 | 115212 | 112731 | 113475 | 1.52E-06 | 1.75E-09 |
| | Upper 95% CI | 20261433 | 910177 | 599007 | 613959 | 629105 | 634054 | 3.15E-06 | 2.26E-06 |
| | | **M_R>AK** | **M_AK>R** | **M_NW>AK** | **M_AK>NW** | **M_R>E** | **M_E>R** | **M_NW>E** | **M_E>NW** |
| | Estimate | 1.27E-06 | 3.35E-07 | 3.24E-06 | 1.63E-07 | 3.26E-08 | 8.29E-09 | 5.13E-07 | 4.83E-06 |
| | Lower 95% CI | 2.92E-09 | 1.81E-09 | 1.87E-06 | 1.95E-09 | 2.10E-09 | 1.50E-09 | 6.66E-10 | 6.15E-10 |
| | Upper 95% CI | 2.80E-06 | 6.53E-06 | 3.51E-06 | 3.74E-06 | 2.35E-06 | 3.55E-06 | 8.34E-05 | 1.08E-04 |
| | | **M_NW>R** | **M_R>NW** | | | | | | |
| | Estimate | 1.66E-07 | 3.70E-06 | | | | | | |
| | Lower 95% CI | 7.47E-10 | 6.91E-10 | | | | | | |
| | Upper 95% CI | 2.73E-05 | 6.95E-05 | | | | | | |

| | | | | | | | | | |
|---|---|---|---|---|---|---|---|---|---|
| Hairy Woodpecker | | **Ne_AK** | **Ne_E** | **Ne_R** | **Ne_NW** | **B-Ne_AK** | **B-Ne_E** | **B-Ne_R** | **B-Ne-NW** |
| | **Estimate** | 7567355 | 18365368 | 47953365 | 5059890 | 804416 | 1486729 | 4693555 | 1515766 |
| | **Lower 95% CI** | 1015730 | 10152829 | 20313563 | 1015965 | 31680 | 1187902 | 12159534 | 22617 |
| | **Upper 95% CI** | 12108980 | 86705380 | 94821663 | 6615990 | 44404284 | 1913030 | 51097323 | 2.3E+07 |
| | | **Ne-AK+E** | **Ne-NW+R** | **Anc_Ne** | **T_div** | **T_div_E** | **T_div_W** | **T_exp_AK** | **T_exp_E** |
| | **Estimate** | 33510699 | 30829288 | 2751382 | 873091 | 425555 | 508965 | 319711 | 347851 |
| | **Lower 95% CI** | 27603525 | 10155085 | 1033504 | 848731 | 272556 | 405623 | 115956 | 115921 |
| | **Upper 95% CI** | 52560389 | 67639132 | 3651731 | 926396 | 486276 | 571448 | 529659 | 633082 |
| | | **T_exp_R** | **T_exp_NW** | **M_E>AK** | **M_AK>E** | **M_R>AK** | **M_AK>R** | **M_NW>AK** | **M_AK>NW** |
| | **Estimate** | 350026 | 335266 | 8.30E-07 | 8.89E-08 | 3.22E-07 | 9.78E-09 | 7.99E-07 | 3.90E-07 |
| | **Lower 95% CI** | 307984 | 117679 | 1.98E-09 | 6.70E-09 | 1.34E-09 | 2.76E-09 | 1.28E-09 | 1.73E-09 |
| | **Upper 95% CI** | 356689 | 429556 | 5.81E-06 | ‡ | 3.55E-06 | ‡ | 4.11E-06 | 3.88E-06 |
| | | **M_R>E** | **M_E>R** | **M_NW>E** | **M_E>NW** | **M_NW>R** | **M_R>NW** | | |
| | **Estimate** | 3.87E-07 | 4.07E-10 | 5.17E-07 | 9.91E-07 | 4.23E-09 | 1.45E-06 | | |
| | **Lower 95% CI** | 1.24E-09 | 3.07E-10 | 1.05E-07 | 1.81E-09 | 2.41E-09 | 2.22E-06 | | |
| | **Upper 95% CI** | ‡ | 5.03E-10 | ‡ | 5.62E-06 | ‡ | 4.02E-06 | | |

Ne_[*pop*]: current $N_e$ in population *pop*; B-Ne_[*pop*]: $N_e$ during bottleneck in population *pop*; Ne_[*pop1*+*pop2*]: $N_e$ in population ancestral to *pop1* and *pop2*; Anc_Ne: ancestral $N_e$; T_div: time of divergence (in years); T_div_E: time of divergence of the Eastern clade (in years); T_div_W: time of divergence of the Western clade (in years); T_exp_[*pop*]: time of expansion of population *pop* (in years); M_[*pop1*>*pop1*]: migration rate (in percent of $N_e$) from *pop1* to *pop2*. ‡: Confidence intervals could not be determined because point estimates fell outside the range of values estimated from bootstrap simulations
